# Immuno-electron microscopy localizes *Caenorhabditis elegans* vitellogenins along the classic exocytosis route

**DOI:** 10.1101/2022.06.27.497668

**Authors:** Chao Zhai, Nan Zhang, Xi-Xia Li, Xue-Ke Tan, Fei Sun, Meng-Qiu Dong

## Abstract

Vitellogenins (VITs) are the most abundant proteins in adult hermaphrodite *C. elegans*. VITs are synthesized in the intestine, secreted to the pseudocoelom, matured into yolk proteins (YPs), and finally deposited in oocytes to support embryonic and larval development. How VITs are secreted out of the intestine remains unclear. In this study, we use immuno-electron microscopy (immuno-EM) to characterize the wild-type subcellular structures containing VITs or YPs. In the intestinal cells of young adult worms, we identify VITs along an exocytic pathway consisting of the rough ER, the Golgi, and the lipid bilayer bounded vesicles, which we call intestinal vitellogenin vesicles (VVs). This suggests that the classic exocytotic pathway mediates secretion of VITs from the intestine to the pseudocoelom. We also show that pseudocoelomic yolk patches (PYPs) are membrane-less and amorphous. The different VITs/YPs are packed as a mixture into the above structures. The size of VVs can vary with the VIT levels and the age of the worm. On adult day 2 (AD 2), intestinal VVs (∼200 nm in diameter) are smaller than gonadal yolk organelles (YOs, ∼500 nm in diameter). VVs, PYPs, and YOs share a uniform, medium electron density by conventional EM. The morphological profiles documented in this study can serve as a reference for future studies of VITs/YPs. Surveying the findings from this study and elsewhere, we review in the discussion the post-translational modifications and protein-protein interactions of *C. elegans* VITs/YPs.

## INTRODUCTION

Vitellogenins belong to an ancient family of large lipid transfer proteins (LLTP) that are found in nearly all metazoans (Wu et al., 2013). A subgroup of this protein family evolved into microsomal triglyceride transfer proteins (MTPs), which are present in vertebrates (Anderson et al., 1998). Vitellogenins and MTPs both have a lipid-binding domain named vitellogenin_N, as does apolipoprotein B (apoB), whose dysregulation in old people may cause cardiovascular disease (Borén & Williams, 2016; Olofsson & Borèn, 2005).

Vitellogenins are precursors of yolk proteins (YPs). They are synthesized by oviparous animals in certain somatic cells outside the gonad, e.g., the fat body of insects, the intestine of sea urchin, and the liver of fish, birds, or amphibians. After posttranslationally modified in the endoplasmic reticulum (ER) and the Golgi apparatus, vitellogenins are secreted as lipoprotein complexes, transported to and deposited into oocytes as nutrients for the development of embryos (Du et al., 2017; Li & Zhang, 2017). Many studies have shown that receptor-mediated endocytosis mediates the uptake of vitellogenins into oocytes, but how vitellogenins are secreted is studied less (Grant & Hirsh, 1999; Li & Zhang, 2017; Perez & Lehner, 2019).

The *C. elegans* animal model has six vitellogenin proteins, VIT-1 to VIT-6. All six VIT proteins have the same modular design, consisting of a signal peptide (SP) at the N-terminus, followed by a DUF1943 domain of unknown function, and a cysteine-rich von Willebrand factor type D (VWD) domain, which is illustrated in Fig. 1A using VIT-2 as an example. Analyzing the sequences of the six *C. elegans* vitellogenins (from wormbase.org), we find that VIT-1 and VIT-2, both precursors of the yolk protein YP170B, share 93.9% sequence identity. VIT-3, VIT-4, and VIT-5 are nearly identical precursors of YP170A. It is difficult to separate YP170A and YP170B on SDS-PAGE because they are similar in size, but because they are less than 60% identity in sequences, they are easily distinguishable by protein or peptide sequencing, e.g., through mass spectrometry analysis. VIT-6 is the most divergent member. It shares only ∼30% sequence identity with either VIT-2 or VIT-5, and it goes through one more round of cleavage after it is processed into YP180, which leads to the formation of YP88 and YP115 (Fig. 1B) (Sharrock, 1984). There are two YP complexes, called B dimer and A complex (Sharrock et al., 1990). B dimer consists of two copies of YP170B, or VIT-1/2. A complex consists of one copy each of YP170A (VIT-3/4/5), YP88, and YP115 (Fig. 1C) (Perez & Lehner, 2019; Sharrock et al., 1990). Both complexes contain about 15% lipids by weight (Sharrock et al., 1990).

**Figure 1.**
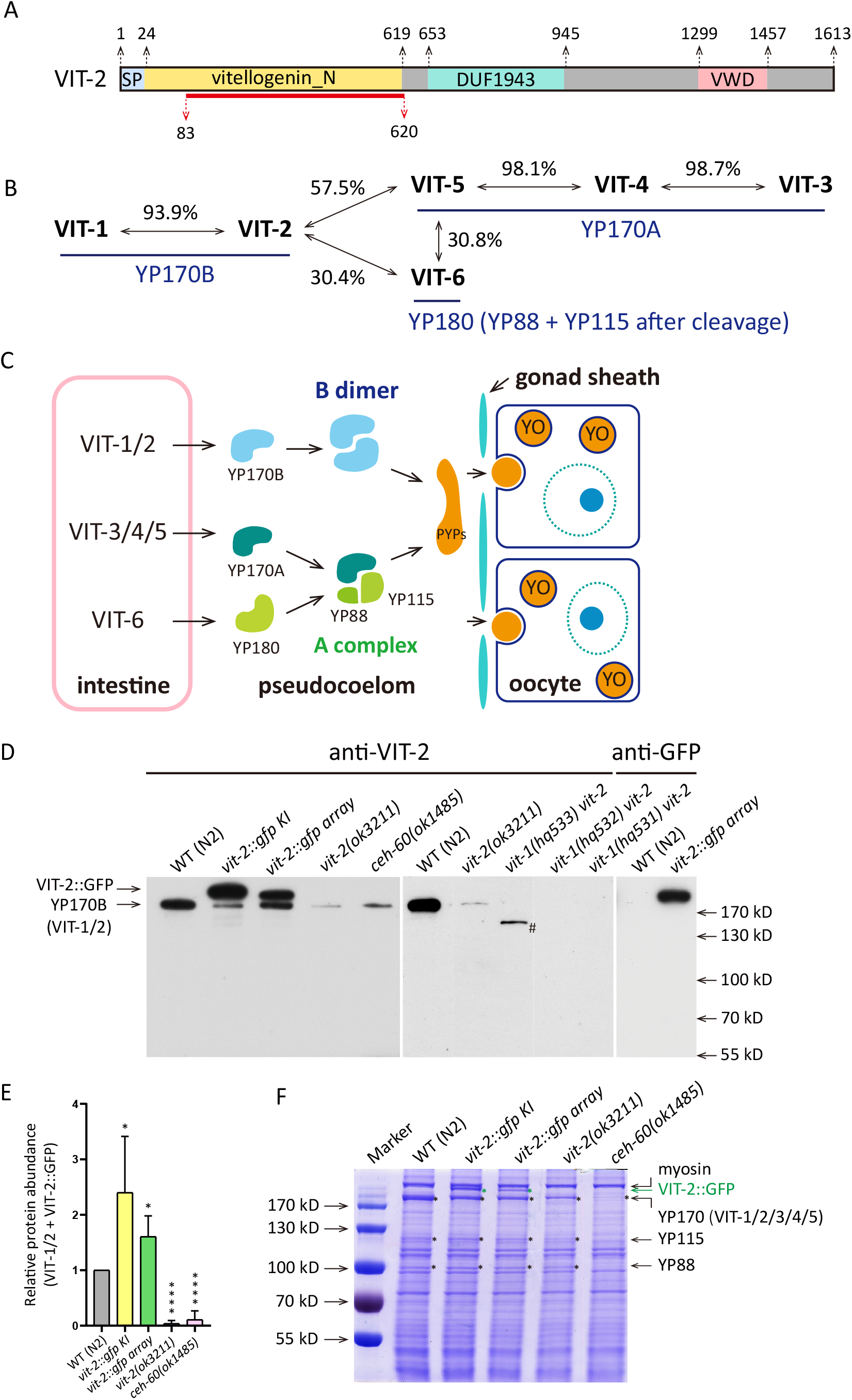
The VIT-2 antibody specifically recognizes YP170B or VIT-1/2. (A) Protein domains of VIT-2. The amino acid positions are indicated above the schematic diagram and the fragment used to immunize rats is indicated below. (B) Protein sequence identity among *C. elegans* vitellogenins, with the names of the corresponding yolk proteins shown below in blue. (C) Formaon and deposition of two types of yolk protein complexes. YO, yolk organelles; PYPs, pseudocoelomic yolk patches. (D) Anti-VIT-2 and an-GFP western blots of whole-worm lysates. # indicates a mutated VIT-1 fragment expressed from hq533. (E) Quantification of the anti-VIT-2 western blot in (D). The intensities of YP170B and VIT-2::GFP, if present, are summed together and normalized against the WT. The bar graph displays the Mean ± STD of eight biological repeats. \**p* < 0.5, \*\*\*\**p* < 0.0001, one-way ANOVA with Tukey’s multiple comparisons test. (F) Coomassie-stained SDS-PAGE of whole-worm lysate to compare the amounts of yolk proteins.

Yolk proteins are the most abundant proteins in adult *C. elegans* hermaphrodites (Sharrock, 1983; Sharrock, 1984; Ezcurra et al., 2018). Vitellogenins are synthesized in the intestine, starting at the time of L4/adult molting (Kimble & Sharrock, 1983). Expression of the *vit* genes is turned on by a homeobox transcription factor (TF) CEH-60 and other TFs (Dowen, 2019; Van de Walle et al., 2019; Van Rompay et al., 2015). After vitellogenins are synthesized and their signal peptides removed, they are secreted by the intestinal cells into the pseudocoelom, which is sometimes referred to as the body cavity. There, they are taken up by developing oocytes through RME-2-mediated endocytosis (Grant & Hirsh, 1999). It is shown that oocytes take up YPs directly from the body cavity through pores in the gonad sheath (Hall et al., 1999). YPs are stored in late-stage oocytes and early embryos in membrane-bound vesicular structures called yolk organelles (YOs), also known as yolk granules, yolk spheres, or yolk platelets (Borgonie et al., 1997; Britton & Murray, 2004; Hall et al., 1999; Paupard et al., 2001). In recent years, studies show that continued production of YPs after reproduction ends causes detrimental aging phenotypes and shortens lifespan (Ezcurra et al., 2018; Kern et al., 2021; Murphy et al., 2003; Seah et al., 2016; Sornda et al., 2019; Wang et al., 2018).

Despite 40 years of research on *C. elegans* vitellogenins, it is not entirely clear how vitellogenins are secreted out of the intestine. The subcellular structures containing VIT/YP are characterized insufficiently in *C. elegans*, especially those located in the intestine. For instance, a literature search finds that two morphologically distinct EM structures in the intestine are both labeled as yolk granules (Lemieux & Ashrafi, 2014; Wolkow & Hall, 2013), and their precise membrane structures are not discernable. This has to do with a lack of systematic and high-resolution immuno-EM analysis of VIT/YP (Britton & Murray, 2004; Hall et al., 1999; Herndon et al., 2002; Paupard et al., 2001).

Using immuno-EM coupled with high-pressure freezing and other microscopy methods, we characterized systematically and quantitatively the vitellogenin- or YP-containing subcellular structures in *C. elegans*. More than validating previously characterized YOs and PYPs with finer resolution, this study identified the precise vitellogenin-containing structures in the intestine, and showed that vitellogenins were secreted via exocytosis from the intestine to pseudocoelom. We clarified that the intestinal vitellogenin vesicles are enclosed by a single lipid bilayer membrane, different from YOs in size, but similar to YOs and PYPs in electron density. This work lays the foundation for further studies on yolk proteins.

## RESULTS

### The anti-VIT-2 antibody specifically recognizes YP170B or VIT-1/2

To document vitellogenin- or YP-containing subcellular structures by immuno-EM, we started by characterizing an anti-VIT-2 antibody. This antibody was generated by immunizing rats with a purified recombinant protein of VIT-2 (amino acids 83-620)::6xHis (Fig. 1A) (Liu et al., 2012). Western blotting indicated that the anti-VIT-2 antibody recognized a single protein band from the lysate of wild-type (WT) worms, which was markedly reduced, but not eliminated, by deletion of either *vit-2* or *ceh-60* (Fig. 1D and E). The *ceh-60* mutant has greatly reduced yolk protein levels (Van de Walle et al., 2019), as confirmed here by SDS-PAGE (Fig. 1F). We reasoned that the remaining signal on the WB likely comes from VIT-1, since VIT-1 and VIT-2 share 93.3% sequence identity across the full length and 96.3% identity across the fragment that was used to generate the antibody. This was verified using three *vit-1 vit-2* double knock-out strains we generated (Fig. 1D). Therefore, this polyclonal antibody recognizes both VIT-2 and VIT-1.

### VIT-2 is found in all VIT- or YP-containing structures in C. elegans

We then asked whether this anti-VIT-2 antibody can label all VIT/YP-containing structures in *C. elegans*. Both VIT-1 and VIT-2 can form the B-dimer complex, while VIT-3/4/5 and VIT-6 form the A complex (Fig. 1C) (Perez & Lehner, 2019; Sharrock et al., 1990). However, it is unclear whether these YP complexes amalgamate together or there are, for example, VIT-1 only or A complex only yolk organelles. Knowing the answer to this question will facilitate interpretation of the immuno-EM data and determine whether all six VITs share the same pathway of transportation from the intestine to oocytes.

Using the CRISPR genome editing method (Dickinson et al., 2013), we engineered three dual fluorescence reporter strains: *hq503(vit-1::mCherry vit-2::gfp), hq485(vit-2::gfp vit-3::mCherry)*, and *hq486(vit-2::gfp; vit-6::mCherry)*. To differentiate the signals of the fluorescent fusion proteins from the auto-fluorescence emitted by gut granules, a type of lysosome-like organelles that are found abundantly in the intestine (Clokey & Jacobson, 1986; Coburn et al., 2013; Coburn & Gems, 2013; Hermann et al., 2005), we acquired epifluorescent images in green, red, and blue channels (Fig. 2-4, Fig. S1, and Fig. S2). The blue fluorescence signals were used to subtract autofluorescent objects. After subtraction, we quantified the colocalization rate of the GFP signal and the mCherry signal in each of the rectangular regions of interest (ROI) distributed in the imaged intestine, pseudocoelom, oocytes, and embryos (10-70 images of 10-31 worms for each category, 3 ROIs per images, 60-200 μm^2^ per ROI.). More details about the quantification analysis can be found in figure S1D-I and the Materials and Method section. On average, ≥ 93% of the VIT-2::GFP signals colocalized with the mCherry signals of VIT-1, -3, and -6 (Fig. 2-4, Fig. S1A-C). These results suggest that A complex and B dimer colocalize, and the B dimer subspecies (VIT-1 homodimer, VIT-2 homodimer, and VIT-1/VIT-2 heterodimer, if there are), also colocalize. Therefore, the transportation route to be mapped using the anti-VIT-1/2 antibody should be an inclusive one for all VITs/YPs.

**Figure 2.**
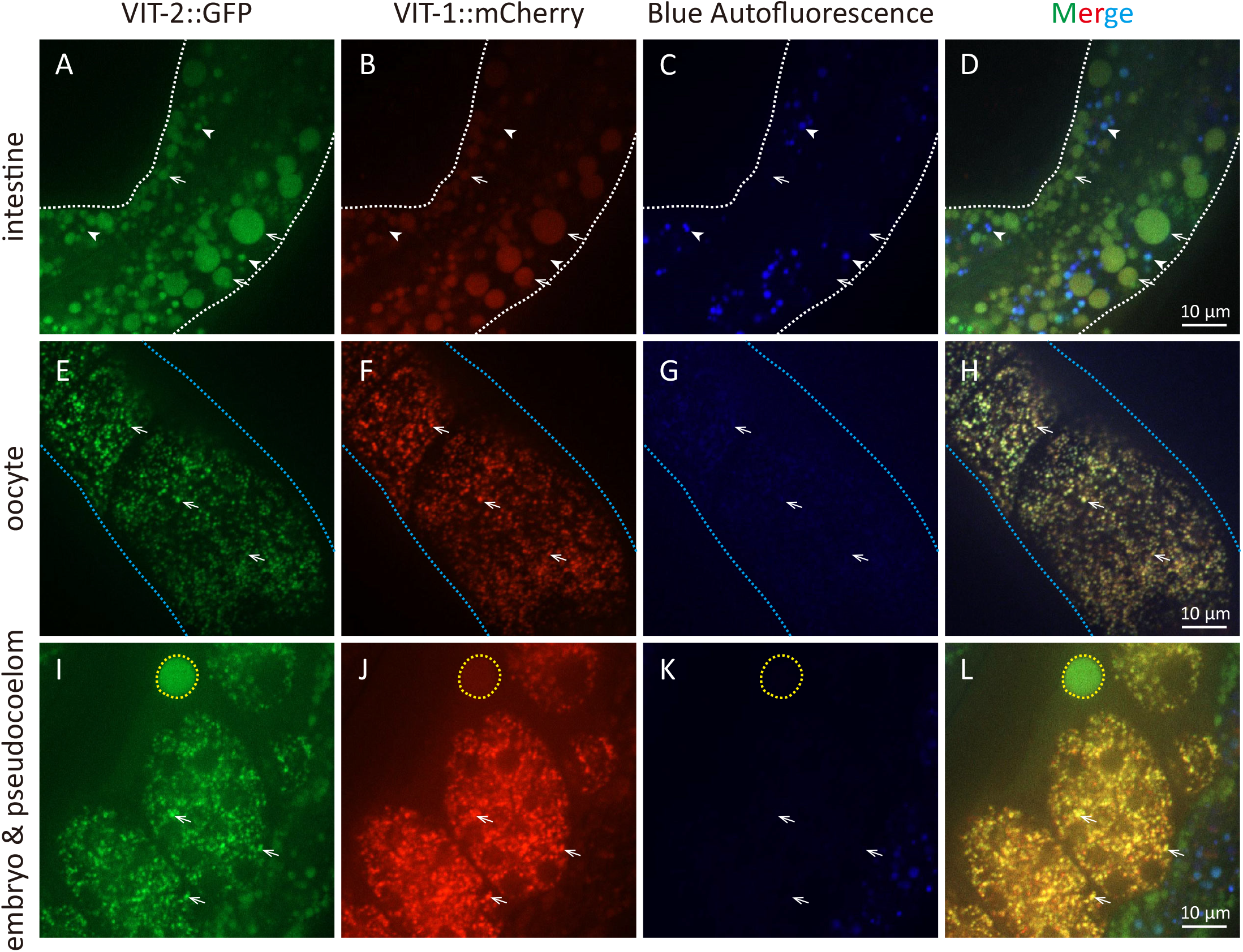
VIT-1::mCherry and VIT-2::GFP colocalize Fluorescent images of *vit-1::mCherry vit-2::gfp* KI worms at AD 2. (A-D) White dots outline the intestine, arrows point to VVs, and arrowheads indicate gut granules, which emit strong green and blue auto-fluorescence. (E-L) Arrows point to YOs in oocytes and embryos. Blue and yellow dots outline the oviduct and a PYP, respectively.

**Figure 3.**
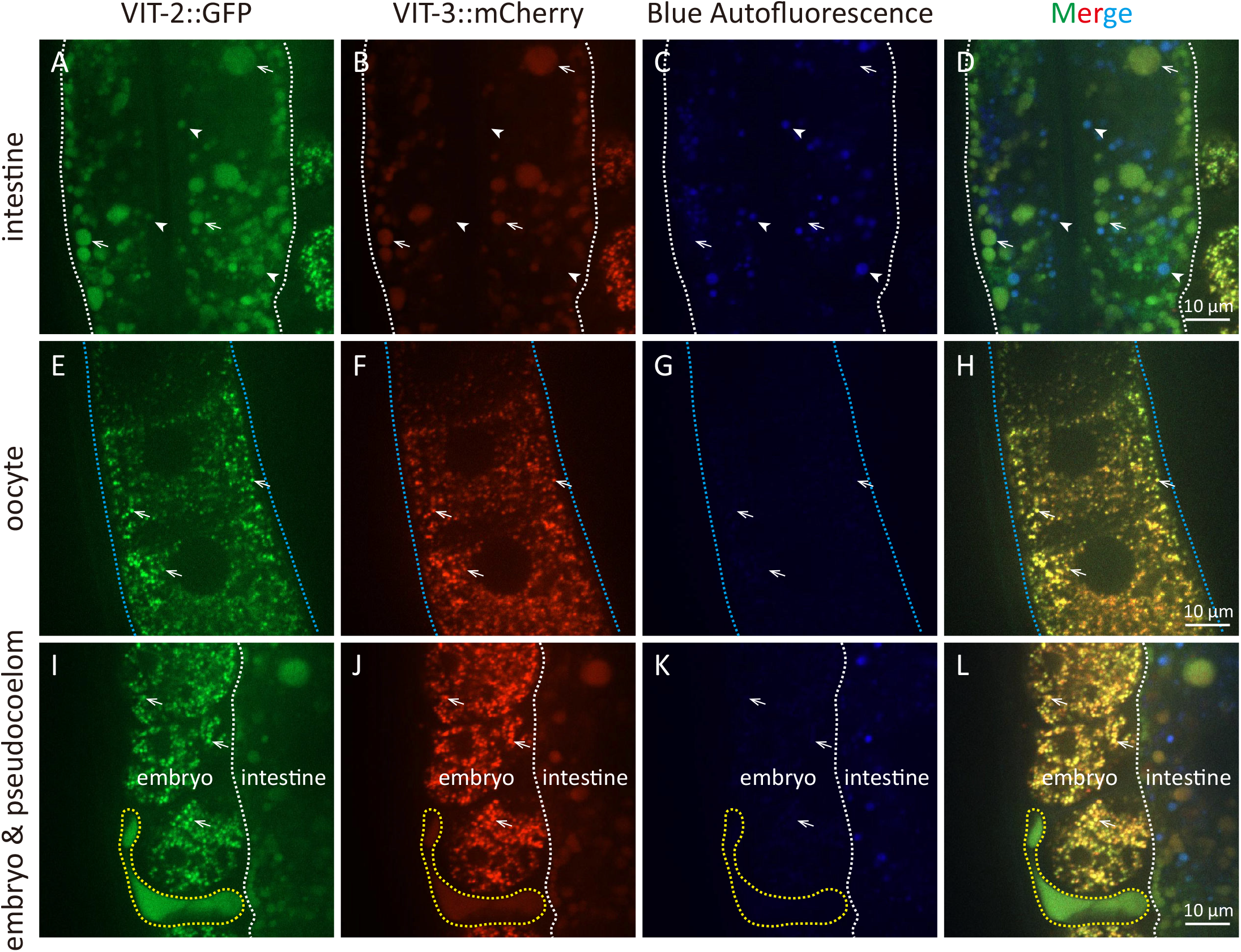
VIT-2::GFP and VIT-3::mCherry colocalize Fluorescent images of *vit-2::gfp vit-3::mCherry* KI worms at AD 2. VVs in the intestine (A-D), YOs in oocytes (E-H), and YOs in embryos (I-L) are indicated by arrows. White, blue, and yellow dots outline the intestine (A-D), the oviduct (E-H), and a PYP (I-L), respectively. The auto-fluorescent gut granules are indicated by arrowheads (A-D).

**Figure 4.**
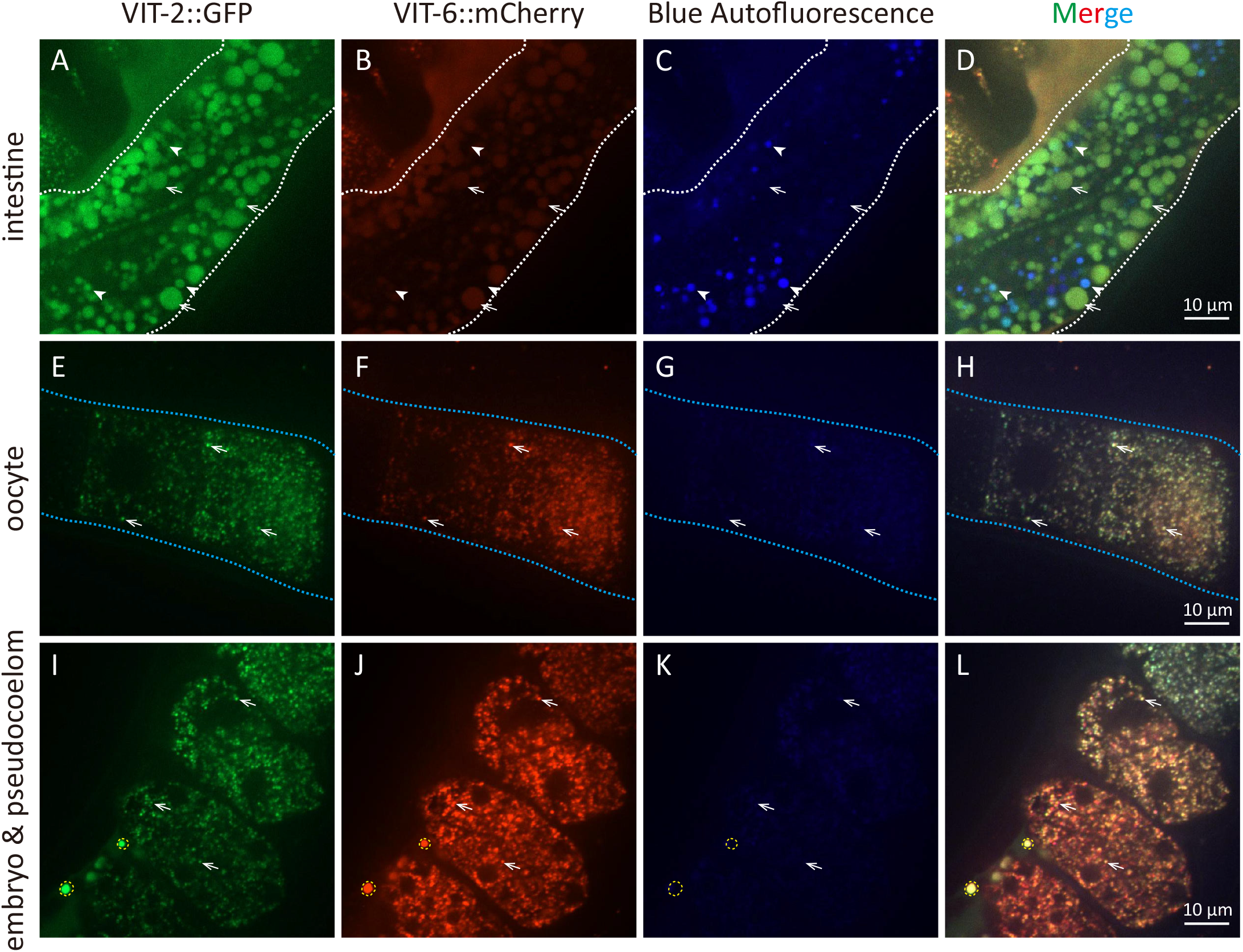
VIT-2::GFP and VIT-6::mCherry colocalize Fluorescent images of *vit-2::gfp; vit-6::mCherry* KI worms at AD 2. VVs in the intestine (A-D), YOs in oocytes (E-H), and YOs in embryos (I-L) are indicated by arrows. White, blue, and yellow dots outline the intestine (A-D), the oviduct (E-H), and a PYP (I-L), respectively. The auto-fluorescent gut granules are indicated by arrowheads (A-D).

### Vitellogenin vesicles in the intestine are smaller than gonadal yolk organelles

Next, we performed immuno-EM analysis. In adult day 2 (AD 2) WT hermaphrodites, the anti-VIT-1/2 antibody targeted gold particles to yolk organelles in embryos and oocytes (Fig. 5A-F), which have been well characterized (Britton & Murray, 2004; Hall et al., 1999). Consistently with previous reports, the observed gonadal YOs were vesicular structures of ∼0.5 μm in diameter (Fig. 5B, F, and I). Since we used high-pressure freezing to prepare samples, which preserved membrane structures well, we were able to see the lipid bilayer membrane of YOs (Fig. 5B).

**Figure 5.**
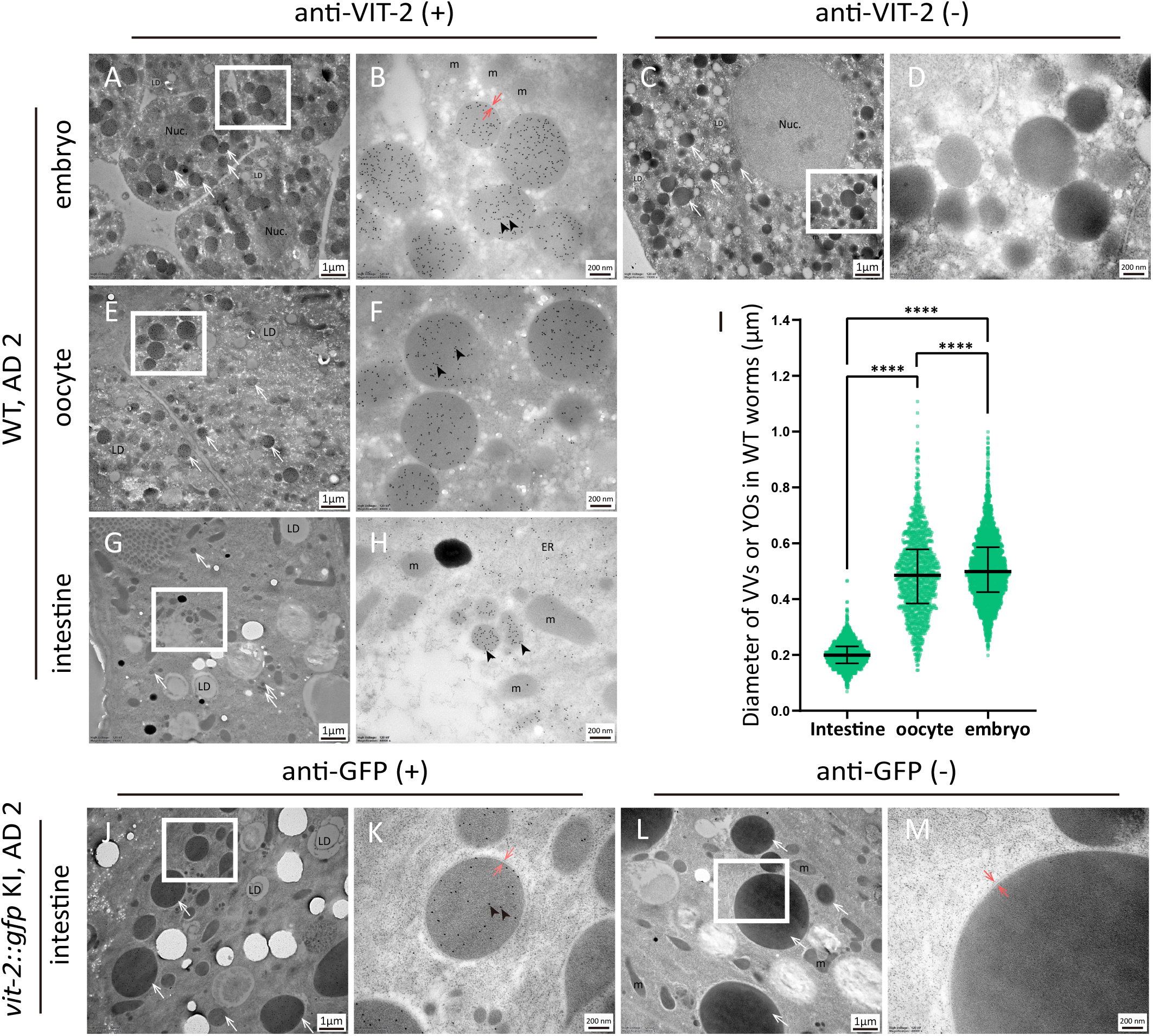
Morphology of intestinal vitellogenin vesicles (VVs) and gonadal yolk organelles (YOs) in young adult *C. elegans* (A-I) Immuno-EM of WT worms on AD 2. (A-B) Micrographs showing anti-VIT-2 immuno-gold labeling of YOs in unlaid embryos in the uterus, (C-D) negative control of (A-B), in which samples were processed without incubation with the anti-VIT-2 antibody. (E-H) Micrographs showing YOs in oocytes (E-F) and VVs in the intestine (G-H) labeled by anti-VIT-2 gold particles. (I) Size analysis of VVs in the intestine, YOs in oocytes, and YOs in unlaid embryos from 217, 58, and 88 micrographs, respectively. For each, the median diameter is indicated along with the interquartile range. \*\*\*\**p* < 0.0001, one-way ANOVA with Tukey’s multiple comparisons test. (J-M) AD 2 worms expressing VIT-2::GFP from a knock-in allele. An-GFP antibody targeted immuno-gold particles to intestinal VVs (J-K). There was no gold particle labeling in the an-GFP (-) negative control (L-M). Black arrowheads point to the 10-nm (A-B, E-H) or 15-nm (J-K) gold particles and white arrows point to gonadal YOs (A-F) or intestinal VVs (G-H, J, L). Paired red arrows indicate the lipid bilayer membrane. The regions framed by white rectangles are magnified and shown on the right. Nuc.: nucleus; LD: lipid droplet; m: mitochondrion; ER: endoplasmic reticulum.

In addition to YOs in the gonad, anti-VIT-1/2 immuno-gold particles labeled similar but much smaller vesicular structures inside the intestine (Fig. 5G-H). The average diameter of these VIT-containing structures was only 0.2 μm (Fig. 5I). As these intestinal structures should contain vitellogenins, not mature YPs (Sharrock, 1984), we termed them vitellogenin vesicles (VVs). VVs were also identified by anti-GFP immuno-EM in the *vit-2::gfp* KI worms (Fig. 5J-M).

### Classic exocytosis mediates vitellogenin secretion from the intestine to the pseudocoelom

In intestinal cells where VIT proteins are synthesized, we observed anti-VIT-1/2 gold particles on the rough ER and the Golgi apparatus (Fig. 6A-D). This is consistent with vitellogenins as secreted proteins, each containing a signal peptide for docking onto the ER. Quantitation of the micrographs showed that the density of the immuno-gold particles in areas of stacked ERs is significantly higher than those that do not contain vitellogenins (*e*.*g*., mitochondria, lipid droplets) (Fig. 6E). This analysis also indicated that the density of gold particles in the Golgi and VVs is about nine times higher than that in the ER, suggesting higher concentrations of vitellogenins in the Golgi and VVs. Similar gold particle densities are found on the Golgi, VVs, and gonadal YOs (Fig. 6E).

**Figure 6.**
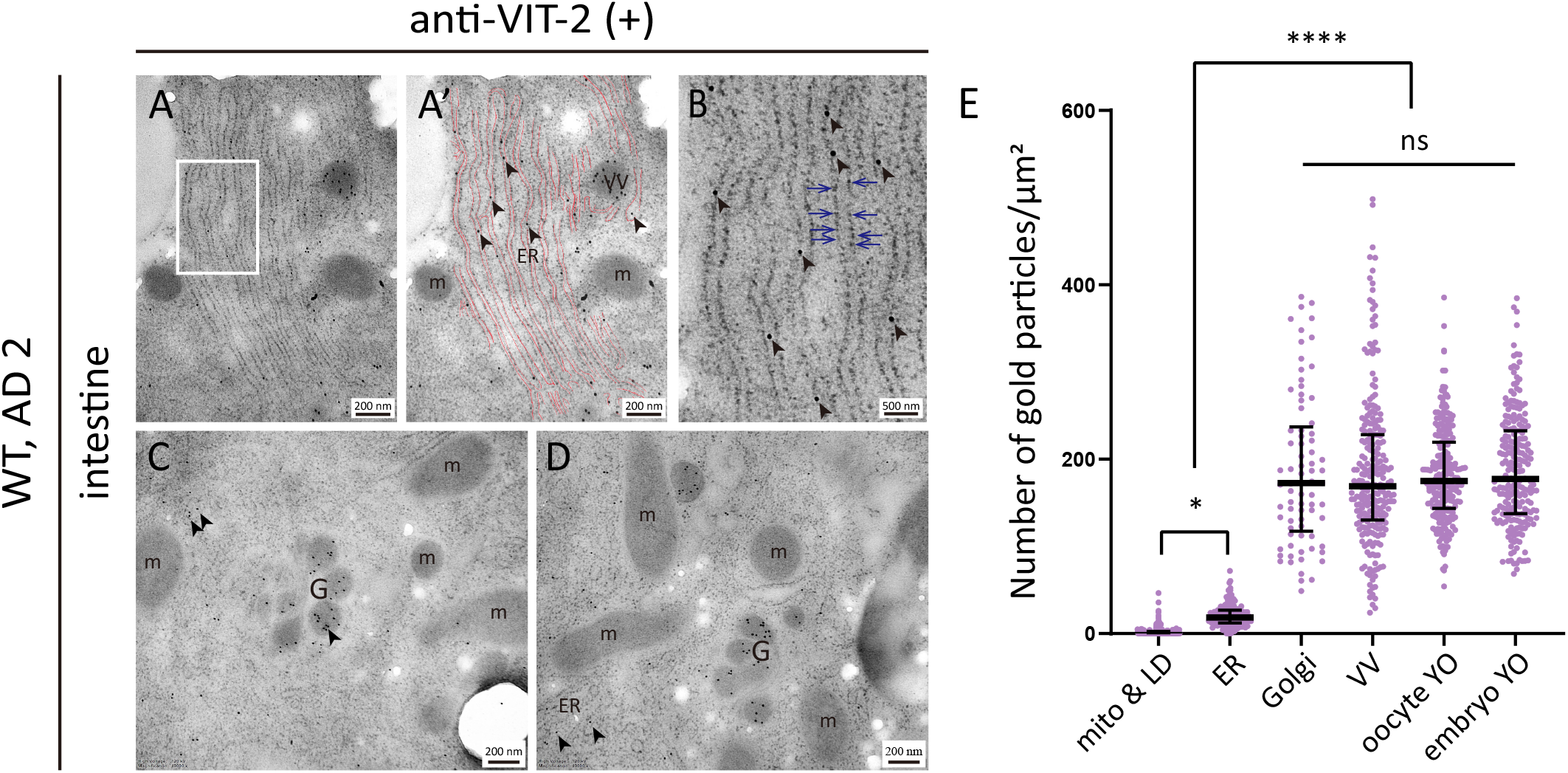
Anti-VIT-2 immuno-gold labeling of the ER and the Golgi apparatus of intestinal cells (A-B) Micrographs showing gold particles attached to the ER. (A) An area of stacked ER labeled by gold particles. (A’) The membrane of ER in (A) is traced by red lines. (B) Magnified view of the boxed region in (A). (C-D) Micrographs of the Golgi labeled by gold particles. ER: endoplasmic reticulum; m: mitochondrion; VV: vitellogenin vesicle; G: the Golgi apparatus. Black arrowheads point to gold particles, blue arrows point to the ribosomal particles attached to the ER. (E) The density of gold particles on different subcellular structures. Measurements were based on 39, 39, 25, 44, 26, and 23 micrographs for mitochondrion and lipid droplet (mito & LD), ER, Golgi, VV, oocyte YO, and embryo YO. The median value and the interquartile range are indicated. \**p* < 0.05; \*\*\*\**p* < 0.0001; ns, not significant; one-way ANOVA with Tukey’s multiple comparisons test.

How *C. elegans* vitellogenins are secreted from the intestine to the pseudocoelom is a topic hardly discussed in the literature. One study on *sft-4*, which encodes an ER exit site protein, came close to but did not go into it (Saegusa et al., 2018). The ER exit site is where secretory proteins are packaged into the coat protein complex II (COPII)-coated transport vesicles, which then bud off and move to the Golgi (Bruce et al., 2002). Knockdown of *sft-4* leads to intestinal accumulation of VIT-2::GFP in the ER lumen (Saegusa et al., 2018). This is consistent with our observation of VIT-1/2 on the rough ER, the Golgi, and VVs by immuno-EM (Fig. 6). All the evidence above points to a classic secretory pathway from the ER to the Golgi, then to the exocytic vesicles, and lastly to the extracellular space after the vesicles fuse with the plasma membrane (PM) (Gomez-Navarro & Miller, 2016). Indeed, we detected yolk pit structures along the basal membrane of intestinal cells (Fig. 7A-B). These yolk pit structures captured a stage right after vesicle-PM fusion when the content of the vesicle is being emptied out. We verified that the yolk substance emptied to the pseudocoelom is not bound by a lipid membrane (Fig. 7C). This is consistent with the varying shape of VIT-positive regions in the pseudocoelom (Fig. 2-4, panels I-L). Yolk substance distributed widely in the pseudocoelom in older worms (AD 6) (Fig. 7C), consistent with the earlier finding (Herndon et al., 2002). Likewise, the yolk pit structures had been observed (Hall et al., 1999), but without the morphological details in membrane structures. It bears emphasis that these VV-PM fusion events are also observable by light microscopy in worms expressing VIT-2::GFP (Fig. 7D-F). Taken together, our data are consistent with the idea that vitellogenins are secreted out of the intestinal cells via a classic exocytosis route (Fig. 7G).

**Figure 7.**
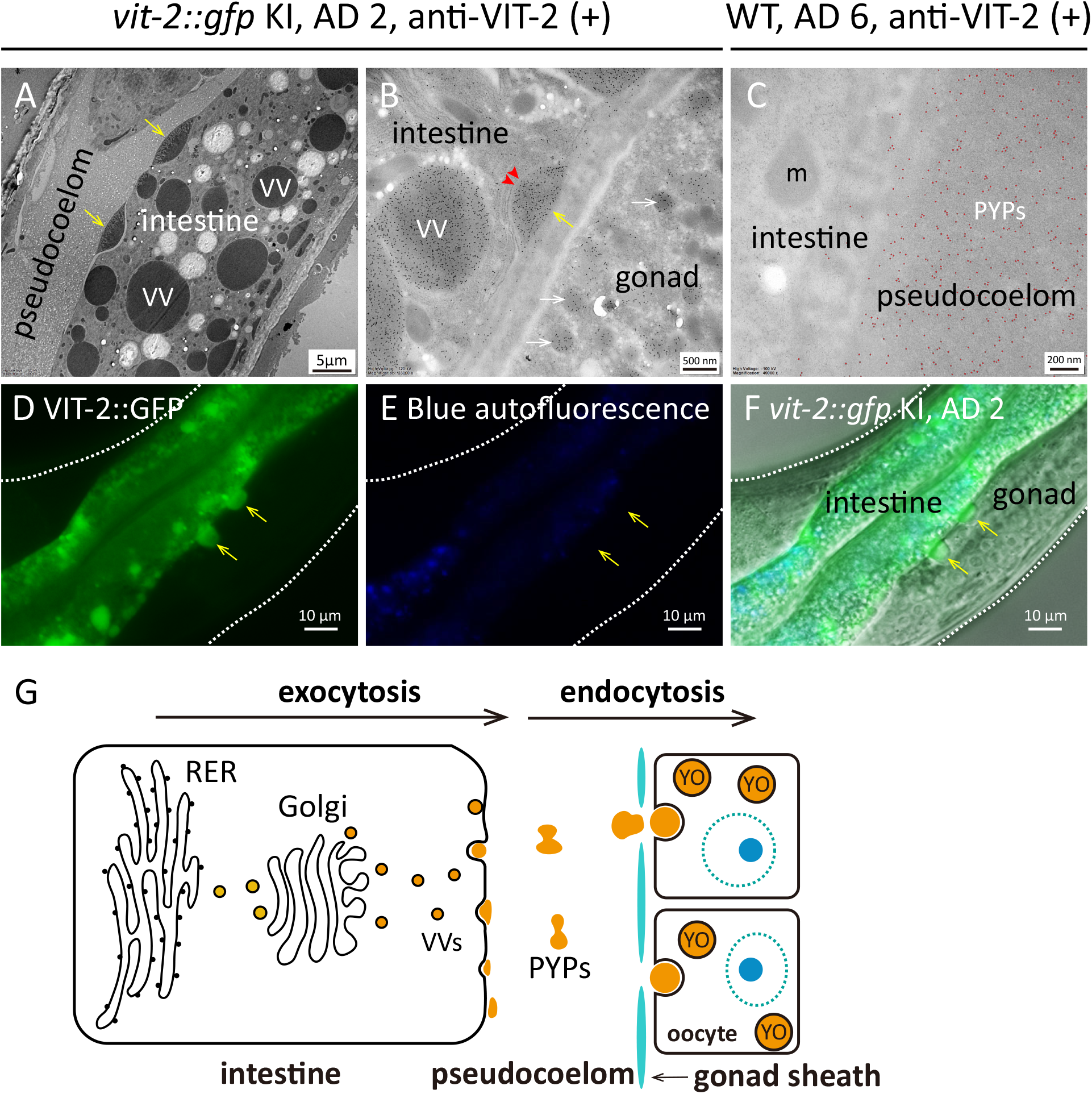
More evidence for exocytosis mediating vitellogenin secretion from the intestine to the pseudocoelom (A-C) Anti-VIT-2 immuno-EM of the *vit-2::gfp* KI worms on AD 2 (A-B) or the WT worms on AD 6 (C). Yellow arrows point to patches of yolk that appear to be at the final stage of exocytosis from the intestine to the pseudocoelom. Red arrowheads point to the basal membrane of the intestine, and white arrows indicate YOs in the gonad. Red dots mark the 10-nm gold particles seen on a patch of yolk in the pseudocoelom. (D-F) Fluorescence imaging captured VIT-2::GFP positive vesicles being released to the pseudocoelom. (G) A model illustrating that vitellogenins are secreted to the pseudocoelom through exocytosis. RER: rough ER; PYPs: pseudocoelomic yolk patches.

### The size of VV but not YO correlated with the total amount of VIT proteins

After we analyzed VIT-containing structures in WT *C. elegans*, we asked how these structures might be affected when VIT proteins are deficient or overly abundant. As verified by SDS-PAGE (Fig. 1F), the yolk protein levels are reduced in the *vit-2* mutant and more so in the *ceh-60* mutant (Van de Walle et al., 2019). Interestingly, the size and the number of intestinal VVs decreased in both mutants (Fig. 8M-N, WT> *vit-2*> *ceh-60*), whereas gonadal YOs were less affected (Fig. 8O). There was no difference in YO size between WT and *vit-2(ok3211)* worms, and the *ceh-60* YOs are smaller by about one third in diameter.

**Figure 8.**
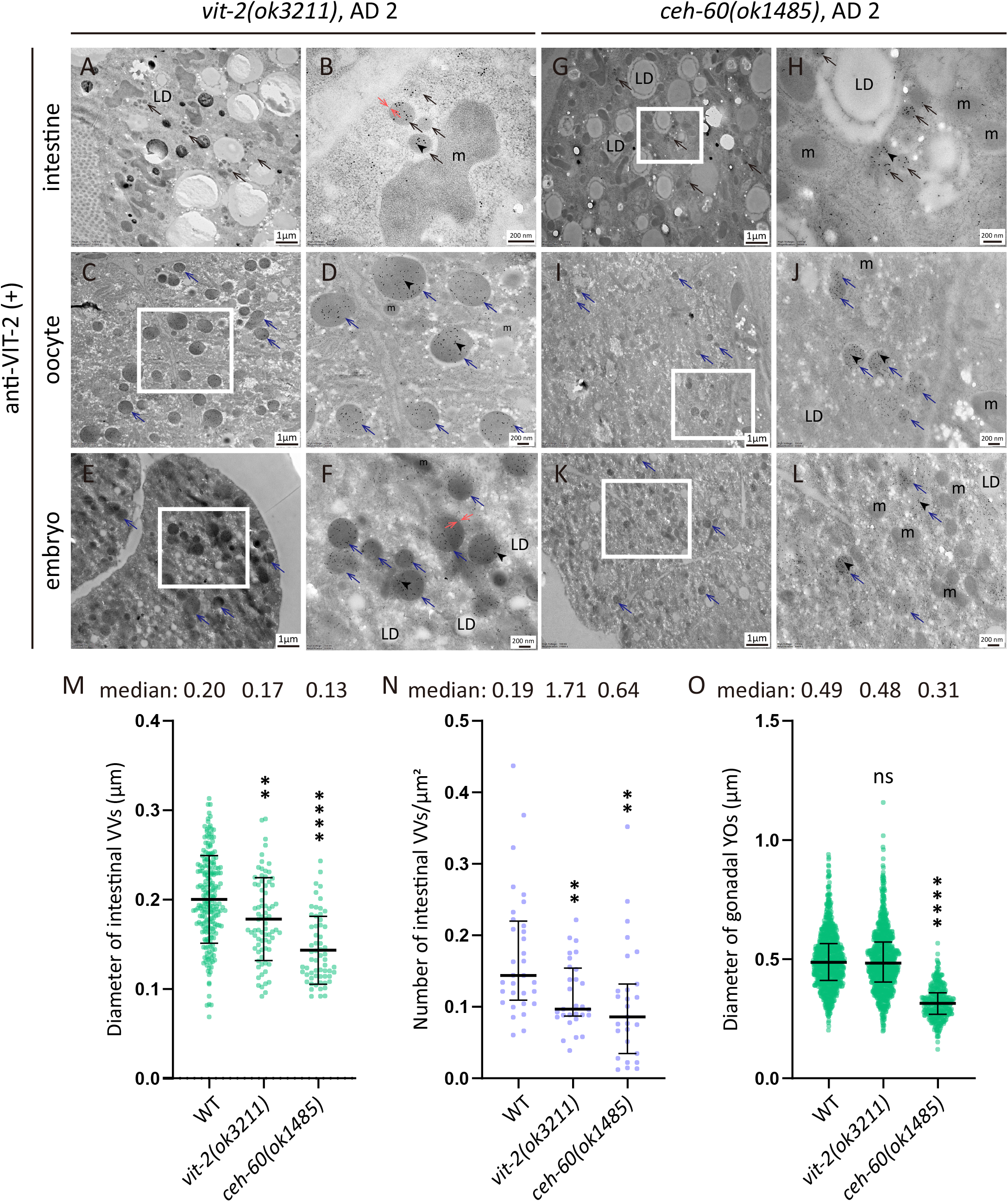
VVs and YOs in VIT-deficient mutants Anti-VIT-2 immuno-EM of the *vit-2(ok3211)* mutant (A-F) or the *ceh-60(ok1485)* mutant (G-L) on AD 2. The boxed regions are magnified and shown on the right. Black arrowheads point to gold particles, black arrows point to intestinal VVs, and blue arrows point to YOs. The lipid bilayer membrane enclosing VVs or YOs is indicated by a pair of red arrows. LD: lipid droplet; m: mitochondrion. (M-O) Quantitative comparison of VVs and YOs between WT and VIT-deficient mutant worms. For WT (N2), *vit-2(ok3211)*, and *ceh-60(ok1485)* worms, 17, 22, and 17 micrographs were quantified for determining the diameter of intestinal VVs (M), 32, 29, and 27 micrographs for the density of intestinal VVs (N), 21, 26, and 25 micrographs for the diameter of gonadal YOs (O), respectively. \*\**p* < 0.01; \*\*\*\**p* < 0.0001; ns, not significant; all compared to WT; one-way ANOVA with Tukey’s multiple comparisons test.

The *C. elegans* strain harboring a *vit-2::gfp* array overexpresses VIT-2::GFP, as does the *vit-2::gfp* KI strain for unknown reason (Fig. 1D-E). In both strains, intestinal VVs enlarged 3-9 times in diameter compared to the WT (Fig. 9A-H and Fig. S3). Consequently, a larger intestinal area (9-13%) was occupied by VVs in these strains compared to only 1% for the WT (Fig. 9I). However, an overabundance of VIT-2::GFP did not increase the size of gonadal YOs. Rather, gonadal YOs were slightly smaller in worms carrying either the *vit-2::gfp* KI allele or the *vit-2::gfp* array (Fig. 9J).

**Figure 9.**
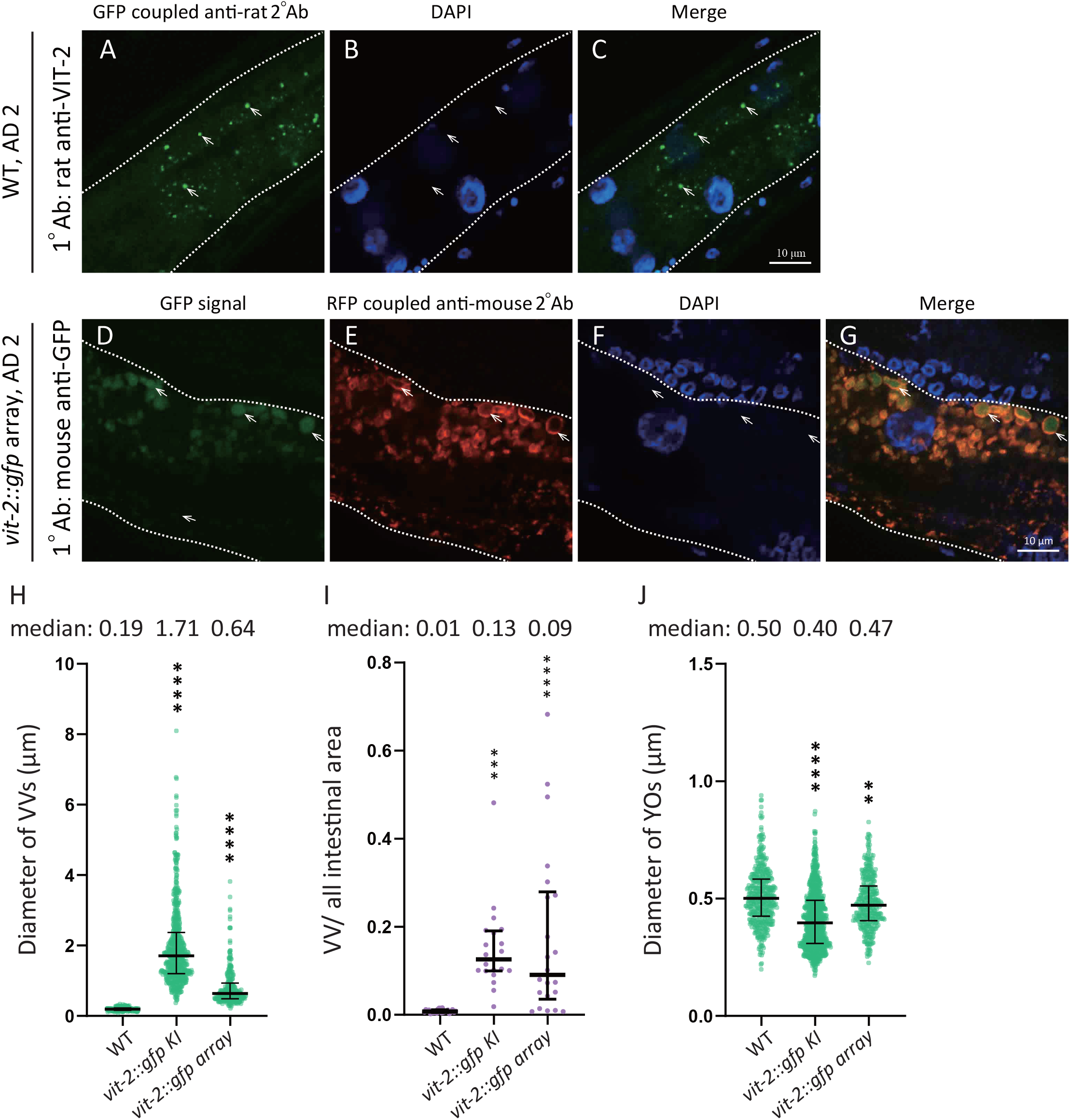
VVs and YOs in worms overexpressing VIT-2::GFP (A-C) Anti-VIT-2 immuno-fluorescence (IF) staining of WT *C. elegans* on AD 2. (D-G) An-GFP IF staining of the VIT-2::GFP overexpressor on AD 2. The GFP tag retained its green fluorescence after IF staining. White dots outline the intestine. (H-J) Quantification of the diameter of intestinal VVs (H), the ratio of VV area over all intestinal area (I), and the diameter of gonadal YOs (I) in WT and two VIT-2::GFP overexpression strains. In the *vit-2::gfp* KI strain, the GFP coding sequence was knocked into the *vit-2* gene locus, which would in theory result in VIT-2::GFP being expressed at or near the endogenous VIT-2 level. However, VIT-2::GFP was somehow expressed at a higher level as shown in Fig. 1D-E. For WT (N2), *vit-2::gfp* KI, and *vit-2::gfp* array strains, 25, 21, and 22 micrographs were quantified for VVs (H), 25, 20, and 22 micrographs were quantified for the ratio of VV area (I), 11, 14, and 11 micrographs were quantified for YOs (J), respectively. Median and interquartile range are indicated. \*\**p* < 0.01; \*\*\*\**p* < 0.0001; all compared to WT; one-way ANOVA with Tukey’s multiple comparisons test.

Taken together, the data above suggest that the size of intestinal VVs but not gonadal YOs correlate positively with the total amount of VIT proteins in the worm.

### Morphology of VIT- or YP-containing structures in conventional electron microscopy

Lastly, to help researchers better identify VVs and YOs using conventional electron micrographs, we followed a previously described approach (Li et al., 2017) and processed two consecutive EM sections differently and then registered and compared the structures on the neighboring sections. Fig. 10A presents an immuno-EM image of the intestine of a WT worm on AD 6, in which a vitellogenin vesicle is labeled by gold particles. Fig. 10B shows the adjacent section, which was not incubated with the antibody; rather, it was stained and collected as if it were a conventional EM section. In other words, Fig. 10B represents a hybrid sample preparation method, in which the samples were processed in the earlier steps for immuno-EM and in the later steps for conventional EM. This allowed us to unambiguously identify which vesicular structures in the intestine are VVs and which are not (Fig. 10C). The typical morphology of AD 2 intestinal VVs, AD 6 gonadal YOs, and AD 6 PYPs are displayed in Fig. 10D-I as a reference. Note that intestinal VVs are larger in older adults (compare panels A-C and D-F of Fig. 10). The morphology of PYPs by conventional EM (Fig. 10C, I) is as same as that by immuno-EM (Fig. 7C), and is consistent with an early report (Herndon et al., 2002).

**Figure 10.**
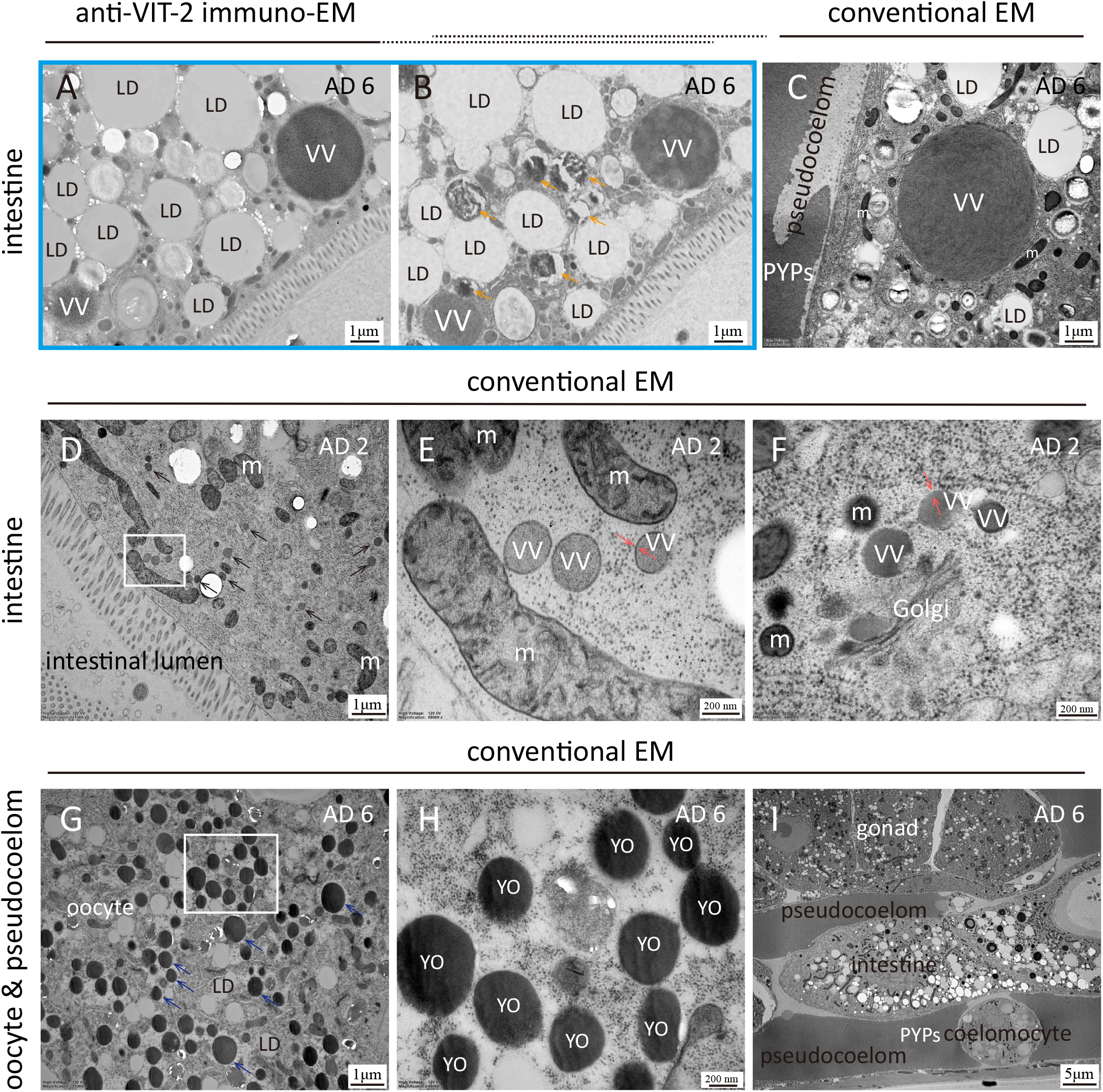
Morphology of intestinal vitellogenin vesicles, gonadal yolk organelles, and pseudocoelomic yolk patches under conventional EM (A-C) Micrographs of intestinal VVs of WT worms on AD 6. (A) Anti-VIT-2 immuno-EM. (B) An adjacent section of the one shown in (A), not incubated with antibodies and colloidal gold particles, but stained with lead citrate and uranyl acetate as in conventional EM, collected on carbon-treated tape, and imaged by scanning EM. (C) conventional EM, imaged by transmission EM (TEM). (D-F) conventional TEM of intestinal VVs of WT worms on AD 2. (G-I) conventional TEM of WT worms on AD 6, highlighting YOs in oocytes (G-H) and PYPs (I). The boxed regions in (D) and (G) are shown at higher magnification in (E) and (H), respectively. The micrographs are marked in the same way as in Fig. 8, except for the orange arrows that depict the μm-sized dark inclusion bodies in Fig. 10B.

## DISCUSSION

### Maturation of YP complexes along the exocytic pathway

The immuno-EM images in this study revealed the presence of vitellogenins in the intestinal cells along the exocytic route, from the rough ER to the Golgi and the vesicles, and right after vesicle-PM fusion. Our finding that secretion of VITs is mediated by classic exocytosis ties together fragments of evidence from earlier studies, analysis of protein sequence features, and detected post-translational modifications.

It is predicted that all six VITs likely have a signal peptide and should direct their synthesis on the rough ER (Teufel et al., 2022). In line with this prediction are several lines of experimental evidence. First, VIT-2::GFP accumulates in the ER when the ER exit site is compromised (Saegusa et al., 2018). Second, YPs are strong Concanavalin A (Con A) binders (Sharrock, 1983), which suggests that YPs likely have high-mannose N-glycans, which are added to proteins in the ER (Roth et al., 2012). Third, YPs have intra- and inter-chain disulfide bonds (Sharrock et al., 1990), which are normally formed in the ER (Fu et al., 2020). Lastly, YP complexes contain 15% lipids by weight (Sharrock et al., 1990), including 8.5% phospholipids, ∼3% TAG, and ∼3% other lipids. Although lipidation of *C. elegans* VITs has not been studied. Lipidation of human apolipoprotein B-100 (ApoB-100) occurs in the lumen of rough ER, associated with microsomal triglyceride transfer protein (MTP) (Hussain et al., 2003). Both ApoB-100 and MTP are homologs of *C. elegans* VIT-1 to VIT-6.

Vitellogenins of non-mammalian vertebrates are heavily phosphorylated, which could be a way of delivering phosphate to oocytes. Phosphorylation is concentrated in a small region called phosvitin, which is not found in *C. elegans* VITs. We looked up a recently published phosphoproteomics dataset and found that each of the six *C. elegans* VITs are phosphorylated at multiple sites (Table S1) (Li et al., 2021). For VIT-1/2/3/4/5, phosphorylation is concentrated near the C-terminus. The phosphosites of VIT-6 are concentrated in the middle region and near the C-terminus.

To sum up, it is most likely that *C. elegans* vitellogenins are large secreted lipoglycophosphoproteins, like their vertebrate counterparts.

### VITs and YP complexes are intermingled in VVs and YOs

Colocalization of VIT-2::GFP and mCherry-tagged VIT-1, VIT-3, and VIT-6 suggests that B dimer (VIT-1/2) and A complex (VIT-3/4/5 and VIT-6) are packed as a mixture into VVs and YOs. The existence of a VIT-1/VIT-2 heterodimer is supported by a previously reported unbiased chemical cross-linking experiment using *C. elegans* protein samples (Table S2) (Tan et al., 2016). From this dataset we found five cross-links between VIT-1 and VIT-2 after applying a stringent filter (#spectra ≥3, best E-val < 1.0 × 10^−7^). For A-complex, we found five cross-links between VIT-3/4/5 and VIT-6, and one cross-link between VIT-3/4 and VIT-6. There was also one cross-link between B dimer and A complex (VIT-2(K972)–VIT-3/4/5(K1201)), suggesting that B dimer and A complex may form a higher-level complex.

mCherry-tagged VIT-1/3/6 appears brighter in developed embryos than in the intestine and oocytes, and the opposite may be said about VIT-2::GFP (see Fig. S4B-C as an example). As YOs are latent lysosomes (Fagotto, 1995), the intensity difference of mCherry and GFP in the developed embryos likely results from acidification of YOs, which activates the lysosomal hydrolases to break down YPs to support embryonic development. Under the acidic pH of the lysosome, GFP loses but mCherry retains fluorescence (Chen et al., 2017). The intensity difference of mCherry and GFP in the intestine or oocytes may be related to the different expression levels of VIT-2 and VIT-1/3/6, at least in part. Mass spectrometry data suggest that VIT-2 and VIT-6 are expressed at higher levels than VIT-1 and VIT-3 in WT worms, as the integrated whole-organism protein abundance values for VIT-1, -2, -3, and -6 are 926, 1555, 838, and 1632 parts per million, respectively (cite pax-db.org). There are other possible reasons. Conditional increase or decrease in the intensity of fluorescent proteins was reported before (Doherty et al., 2010).

### Size control of VVs and YOs

One interesting observation from this study is that intestinal VVs and gonadal YOs seem to have different size control mechanisms. The size of VVs is sensitive to VIT production, changing from 0.13 μm (65% of WT) to 1.71 μm (900% of WT) in diameter, from YP deficient to overproduction strains (Fig. 8-9). In contrast, the size of YOs changes little (0.31-0.48 μm in diameter) and overproduction of YPs does not make bigger YOs (Fig. 8-9). It is suggested that the sheath pore, typically 100-300 nm in diameter, may function as a “screen stencil” (Hall et al., 1999). This hypothesis could explain why YOs are not bigger when YPs are in excess.

### Presence of YOs in early-stage oocytes, albeit fewer than in late-stage oocytes

*C. elegans* hermaphrodites on AD 2 have 8-10 oocytes in each proximal gonad and they are numbered -1, -2, -3, and so on from the one that is most proximal to the spermatheca. A recent review states that yolk in *C. elegans* is loaded into the three most proximal oocytes (Perez & Lehner, 2019), but a close examination of the imaging data (Grant & Hirsh, 1999; Perez & Lehner, 2019) finds that VIT-2::GFP is present in -4, -5, and even earlier-stage oocytes, albeit at an increasingly lower level. This is consistent with what we see by immuno-EM and confocal fluorescent microscopy (Fig. S4). Scanning EM shows that three pairs of sheath cells (No. 3-5) are covered with pores, and these sheath cells wrap around -1 to -6 oocytes (Hall et al., 1999; Lints, 2009). Theoretically, -1 to -6 oocytes can take up yolk directly from the pseudocoelom, but -1 oocyte contains more YOs than -2 oocyte, and -2 more than -3, and so on. That mature oocytes have more time to accumulate yolk is one possible explanation for the formation of this YO gradient. Another explanation is provided by a study on RME-2, the yolk protein receptor. Although oocytes start to express RME-2 from an early stage, the several most proximal ones have more YP-binding extracellular domain exposed on the cell surface (Grant & Hirsh, 1999).

### Are yolk proteins dispensable for embryogenesis?

In *ceh-60* and *vrp-1* mutant worms, the amount of YPs is much reduced, yet the brood size is normal and embryo development seems unaffected (Dowen, 2019; Van Rompay et al., 2015). RNAi of *vit* genes has no effect on fertility, either (Ezcurra et al., 2018). Consequently, the notion that yolk proteins are required for *C. elegans* embryogenesis is challenged (Perez & Lehner, 2019; Van Rompay et al., 2015). Here, immuno-EM readily detected VVs and YOs in the *ceh-60* mutant, albeit of a smaller size (Fig. 8). It follows that the *ceh-60* oocytes likely have enough residual YPs to support embryogenesis. As none of the existing mutations or RNAi treatments completely abolishes yolk proteins, there is not enough evidence to conclude that *C. elegans* yolk proteins are dispensable for embryogenesis.

### VVs are not gut granules defined by light microscopy, nor the dark inclusion bodies defined by EM

Intestinal cells are filled with vesicles of different types, one of which is autofluorescent gut granules (GGs). GGs contain glycosylated anthranilic acid, which emits intense blue fluorescence under UV light (Clokey & Jacobson, 1986; Coburn et al., 2013; Coburn & Gems, 2013; Hermann et al., 2005). A subset of GGs also contains birefringent materials (Hermann et al., 2005). Here we show that VVs (i.e., intestinal yolk granules) contain neither birefringent materials (Fig. S2) nor blue fluorescent materials (Fig. 2-4, Fig. S2). In other words, VVs and GGs are distinct from each other.

Prior to this study, VVs were thought to be similar in size to the well-characterized YOs in oocytes and embryos. Our immuno-EM analysis found that in WT AD2 adults, VVs are of a medium electron density and ∼200 nm in diameter, much smaller than gonadal YOs (∼500 nm). Only in worms of high VIT levels or an older age (e.g., AD 6) did we find μm-sized VVs (Fig. 5J-M and Fig. 10A-C). In two earlier conventional EM studies, the vesicles labeled as “intestinal yolk granules” were almost certainly mis-identified. In one, the “yolk granule” label was attached to μm-sized dark inclusion bodies with or without a nested concentric ring of pitch-blackness in worms at the L2 or dauer larval stage (Wolkow & Hall, 2013), when *vit* genes are not yet turned on (Kimble & Sharrock, 1983). In the other, vesicles with a nested concentric ring of pitch-blackness were labeled tentatively as “maturing yolk?” (Lemieux & Ashrafi, 2014).

### The body wall muscle does not contain VITs in the young WT worms

A recent study reported that body wall muscles (BWM) of young adult worms can synthesize VIT-2, and transport it to oocytes via VIT-2-containing exophers, to promote the development of offspring (Turek et al., 2021). However, our anti-VIT-1/2 immuno-EM analysis did not find evidence for this claim. Neither the body wall muscle nor the pharyngeal muscle were labeled by immuno-gold particles above the background level (Fig. S5). An earlier study has used a sensitive [^35^S]-methionine incorporation method to label newly synthesized proteins and found that only the dissected intestine synthesized yolk proteins, the gonad and the body wall, which contained the BWM, hypodermis and cuticle, did not (Kimble & Sharrock, 1983).

## MATERIALS AND METHODS

### VIT sequences alignment

Download VIT sequences from WormBase, then upload every two sequences of them on the web tool Pairwise Sequence Alignment (https://www.ebi.ac.uk/Tools/psa/emboss_needle/). The percentage identities between every two VITs (Fig. 1B) were calculated via the EMBOSS Needle method, which creates an optimal global alignment of two sequences using the Needle-man-Wunsch algorithm.

### Worm Culture and Strains

*C. elegans* was fed with *E. coli* OP50 on nematode growth medium (NGM) plates and cultured at 20 °C. To produce synchronized cohorts of worms, 25 gravid hermaphrodites were put on a plate and allowed to lay eggs for 4 hours before being taken away. Worms within 24 hours after reaching sexual maturity are regarded as one day old. In this study, eleven strains were used, including wild type (N2), MQD1052 *vit-2(bIs1 [vit-2::gfp + rol-6(su1006)]) X*, BCN9071 *vit-2(crg9070[vit-2::gfp]) X*, MQD2798 *vit-1, vit-2 (hq503[vit-1::mCherry vit-2::gfp]) X*, MQD2775 *vit-2, vit-3 (hq485[vit-2::gfp vit-3::mCherry]) X*, MQD2774 *vit-2, vit-6 (hq486[vit-2::gfp; vit-6::mCherry]) X; IV*, RB2365 *vit-2(ok3211) X*, MQD2883 *vit-1(hq531) vit-2(ok3211)*, MQD2884 *vit-1(hq532) vit-2(ok3211)*, MQD2885 *vit-1(hq533) vit-2(ok3211)*, VC988 *ceh-60(ok1485) X*.

### Worm Strain Construction

The CRISPR method for editing the *C. elegans* genome has been fully developed and described before (Dickinson et al., 2013). We thus used CRISPR and separately knocked *mCherry* into the C terminus of *vit-1, vit-3*, and *vit-6* in a BCN9071 *vit-2(crg9070[vit-2::gfp])* background, and knocked out *vit-1* in an RB2365 *vit-2(ok3211)* background. The pDD162 plasmid (Addgene, Cat. #47549) of Cas9-sgRNA with no target sequence is available as a commercial product. Two target sequences of sgRNA were designed for every knock-in or knock-out strain. The eight target sequences are as follows: *vit-1::mCherry* KI: CGCTTATTAATTCATAAGCT, TATTAATTCATAAGCTCGGC; *vit-3::mCherry KI*: GGGATGTTGGTATTAACAGC, GGATGTTGGTATTAACAGCT; *vit-6::mCherry* KI: TTCACAATCATACACATC GT, GACTGTAGAAGTGAACTCTG; and for *vit-1* KO: TTGGGTATCGACTGGGAGGA, GAGTCCACAACTGT TGTCCG.

Homologous repair templates for *mCherry* KI strains were inserted in the vector 95.77-M5. Sequences of the homologous templates are shown in supplementary files (.dna format file S1-3). The DNA sequences of *vit-1* mutants are shown in supplementary files (.dna format file S4-6). The pRF4 plasmid, which encodes ROL-6 as a marker for selection, was co-injected. The solution for microinjection contained two Cas9-sgRNA plasmids, the pRF4 plasmid, and, if necessary, the plasmid containing homologous repair templates. The final concentration of each plasmid was 50 ng/μl.

Primers designed for genotyping are as follows: for the *vit-1::mCherry* KI stain, WT forward primer, TGATCGCGTCGTCGTTCAACTCAC, and WT reverse primer, AGGCCGGCCGAGCTTATGAATTAAT AAG; KI forward primer, ATTCGCCGCTCCCCAATCCTCGAG, KI reverse primer, ATGTTATCCTCCTCGCCCTTGCTCAC; WT primers amplify an ∼1,200 bp product, while KI primers produce an ∼1,600 bp fragment. For the *vit-3::mCherry* KI stain, WT forward primer, AAGAAGGTTAACCCAACTGAACTTG, WT reverse primer, ACCCCAGCTGTTAATACCAACATCCC; KI forward primer, AGACGGCGAGTTCATCTACAAGGTG, KI reverse primer, AAATGTGCTCAAATGGTTATGCATTG. WT primers amplify an ∼900 bp product, while KI primers produce an ∼1,500 bp fragment. For the *vit-6::mCherry* KI stain, WT forward primer, ATGTACAAGACTGAGGAAGGACTC, WT reverse primer, TGTGTATGATTGTGAAGAAG AGGTAG; KI forward primer, AGACGGCGAGTTCATCTACAAGGTG, KI reverse primer, ATGGTGCAACG CAGTTTGAATTCTAAC. WT primers amplify an ∼600 bp product, while KI primers produce an ∼1,500 bp fragment. For the *vit-1* KO strain, the forward primer, TACCAACGTGTTGCTATCGTTTGCTC, and the reverse primer, TTGCTCGAAGAGTGGGGTGAACATTCTC. This pair of primers amplifies an ∼800 bp product, then sequence PCR product.

### Antibodies

The rat polyclonal anti-VIT-2 antibody (diluted to 1:100 for immuno-EM labeling and to 1:6,000 for western blot) was kindly provided by Dr. Xiao-Chen Wang (Institute of Biophysics, Chinese Academy of Sciences, Beijing, China). The epitope of the antibody is recombinant protein VIT-2 (83-620 amino acid)::6xHIS. The mouse monoclonal anti-GFP antibody is a commercial product from Roche (lot #11814460001, diluted to 1:50 for immuno-EM labeling and to 1:3,000 for western blot). The rabbit-derived secondary antibody (anti-rat) conjugated with 10-nm colloidal gold (Sigma, lot #SLBZ8963), the goat-derived secondary antibody (anti-mouse) conjugated with 6-nm colloidal gold (AURION, code #106.022), and the donkey derived second antibody (anti-mouse) conjugated with 15-nm colloidal gold (Jackson ImmunoResearch, lot #143577) are available as commercial products. The secondary antibodies used in the western blot assay were goat anti-mouse (Sigma, AP124), and goat anti-rat (CW bio, 01340/40126).

### Western Blot Assay and Coomassie Staining

Forty hermaphrodites at adult day 1 were put into 20 μl M9 buffer and frozen in liquid nitrogen as soon as possible. To homogenize the worms, 5 μl 5x SDS sample buffer was added and samples were boiled at 100°C for 10 min. After that, samples were stirred gently using a pipette, then centrifuged at 4°C with 13,000 rpm for 10 min. The supernatant was loaded on a polyacrylamide gel for electrophoresis (PAGE) (80 V, 40 min, 120 V, 110 min). Proteins in the SDS-PAGE gel were transferred to a PVDF membrane at 100 V for 120 min. The anti-VIT-2 antibody and the anti-GFP antibody were diluted to 1:6,000 and 1:3,000 in blocking solution (5% de-fat milk in TBST) for the western blot assay. The anti-rat and anti-mouse antibodies were diluted to 1:5,000 and 1:3,000 in blocking solution.

After electrophoresis, the polyacrylamide gel was placed in the transfer buffer to remove impurities. The polyacrylamide gel was then stained in Coomassie staining solution (0.1% Coomassie Blue R-250, 50% MeOH, 10% HoAC, 39.9% H_2_O) for 4 h. Wash the gel via the destaining solution (50% MeOH, 10% HoAC, 40% H_2_O) before being photographed.

### High-pressure Freezing and Freeze-substitution

The procedure of high-pressure freezing was explained before (Jiang et al., 2020), but a few conditions were optimized here for the nematode samples. The main steps are shown below. Aluminum carriers (Engineering Office of M. Wohlwend GmbH, lot #241) were filled with cryoprotectant (20% BSA in M9 buffer). Ten worms of an indicated age (AD 2, 6, and 9) were put into a carrier. The carriers were covered with sapphire discs (Engineering Office of M. Wohlwend, 3 mm x 0.16 mm). High-pressure freezing was done using the Wohlwend HPF Compact-01 apparatus.

After high-pressure freezing, every four carriers containing samples were put into a 2 mL polypropylene tube (BIOLOGIX, lot #81-0204) containing 1 ml substitution solution, including 1% paraformaldehyde, 4% ddH_2_O, and 95% acetone. Caps with O-rings (BIOLOGIX, lot #81-0004) were screwed into the polypropylene tubes. All steps described above were conducted under liquid nitrogen. Then, tubes were transferred into a Leica AFS2 freeze-substitution machine for dehydration. The temperature controlling procedure was set as follows: -90°C for 48 h; -90°C to - 60°C for 15 h, increase 2°C per hour; -60°C for 8 h; -60°C to -30°C for 15 h, increase 2°C per hour; - 30°C for 8 h; and -30°C to 4°C for 17 h, increase 2°C per hour. When the temperature reached 4°C, precooled acetone was used to rinse samples four times, for 15 min each time.

### Infiltration and Embedding

After dehydration, samples were infiltrated in LR White Resin (London Resin, code #AGR1281A) for 3 days at 4°C. The worm cakes were taken out from carriers carefully using a pair of needles on a 1 ml syringe under a stereo microscope. Then, every worm cake was transferred into the bottom of a gelatin capsule (Electron Microscopy Sciences, Cat. #70110) that was already filled with 2 drops of LR White Resin containing 0.5% accelerator (London Resin, code #AGR1283). The capsules were filled with embedding resin and covered with caps to avoid contact with air. Polymerization was performed under UV illumination for 24 hours at 4°C. After polymerization, the blocks were stored at -20°C before sectioning.

### Sectioning

Blocks were carefully trimmed with a razor under the stereo microscope to make the sample part protrude. Then, a diamond knife (Diatome, DU4530) was used to produce semithin sections (200 nm thick) on the Leica ultramicrotome (UC6, Leica, Germany). Semithin sections were stained with 1% Toluidine blue to check whether the anatomic position of the region of interest was under the microscope (Chinauop, UB103i). After that, 80-nm ultrathin sections were produced using the diamond knife and collected on nickel grids (Electron Microscopy China, Cat. #AG50), one side of which was coated with the formvar film made from formvar resin (SPI-Chem, CAS #63148-65-2). Meanwhile, ultrathin sections were also collected on a piece of tape with glow discharge treatment (Li et al., 2017). Finally, nickel grids carrying sections were stored in grid boxes for immuno-labeling.

### Immuno-labeling

Immuno-labeling was performed mainly according to Weimer et al. (Weimer, 2006), with some modifications. In brief, nickel grids carrying worm sections were incubated in 0.05 M glycine in 0.1 M PB (pH 7.4) for 15 min to quench unreacted aldehyde groups in samples. Then, grids were blocked with 5% BSA in 0.1 M PB (pH 7.4), for 1 h at room temperature, and rinsed three times with 0.1 M PB (pH 7.4). Then, nickel grids were incubated with the primary antibody diluted in antibody buffer (1% BSA in 0.1 M PB, pH 7.4) for 1 h at room temperature. To eliminate unspecific labeling, nickel grids were rinsed six times with 0.1 M PB (pH 7.4), for 5 minutes each time. Then, they were incubated with the second antibody conjugated with colloidal gold for 1 h at room temperature, followed by washing with 0.1 M PB (pH 7.4) nine times, for 5 minutes each time. The interaction between primary and secondary antibodies was fixed using 2% glutaraldehyde in 0.1 M PB (pH 7.4) for 5 minutes, and nickel grids were rinsed three times with distilled water, for 5 minutes each time.

### Transmission EM and Scanning EM Imaging

After staining with uranium acetate and lead citrate, nickel grids were examined using a Tecnai Spirit 120 microscope (FEI, USA) operating at 120 kV. Images were captured with an MoradeG3 CCD (EMSIS) camera using the RADIUS (EMSIS GmbH) software.

Before scanning EM (SEM) imaging, the tapes carrying sections were adhered to SEM Cylinder Specimen Mounts (Electron Microscopy China, Cat. #DP16232) by carbon conductive double-faced adhesive tape (NISSHIN EM Co. Ltd, Japan), then tapes were stained with uranium acetate. The specimen mounts carrying samples were transferred under the SEM (FEI Helios NanoLab 600i) equipped with a CBS detector. Images were acquired by the software xT microscope control (FEI, version 5.2.2.2898) with the SEM parameters set as 2 kV accelerating voltage, 0.69 nA current, and 5 μs dwell time.

### Confocal Imaging

Fifteen live worms were picked and placed on an agarose pad stuck on a glass slide. Worms were paralyzed with 10 mM levamisole aqueous solution and a coverslip was carefully placed on top. Images were captured using a fluorescent microscope. Pictures in Fig. S4A-C were imaged by SpinSR equipped with a 63x oil-immersion objective (magnification 630x). Images were visualized and processed via the OLYMPUS cellSens Dimension. Pictures in Fig. 7D-F and Fig. S2 were imaged by a ZEISS LSM 880 microscope equipped with a 63x oil-immersion objective (magnification 630x). Pictures were visualized with the software Axio Vision Rel. 4.7. Pictures in Fig. 2-4, Fig. 9A-G, Fig. S1D-G, Fig. S3A-G, and images used for quantification were captured using a spinning-disk microscope (UltraVIEW VOX; PerkinElmer) equipped with a 63x oil-immersion objective (magnification 630x). Pictures were visualized with the Volocity software (PerkinElmer).

### Quantification of colocalization between VITs

The imaging data for quantification were preprocessed using the Volocity^®^ Demo 6.3 software. For each original image, the brightness of the green fluorescence and that of the red were adjusted to similar levels, and at the same time it was made sure that the background remained dark for both the green and the red, such as the images in Fig. S1D-G. The adjustment was applied to the entire image and was typically between 1x and 2.5x for either channel. The preprocessed images were saved as .jpg files and further analyzed using the Image J software. Three rectangular frames of 60-200 μm^2^ were randomly placed on a fluorescent image of the intestine, pseudocoelom, oocytes, or embryos (Fig. S1D-H). Each frame defined a region of interest (ROI). Within an ROI, each non-blue-fluorescent punctum (for the intestine and the pseudocoelom) or pixel (for oocytes and embryos) was classified based on its Hue value as GFP only (71-90), mCherry only (1-14), or GFP+mCherry (i.e., colocalized, 15-70). A total of 10-70 images of 10-31 worms were quantified for each category. The ratio of colocalization was calculated as the number of colocalized puncta or pixels divided by the total number of vesicles or pixels for each ROI. The hue value 15-70 was used to define the GFP+mCherry ROIs to accommodate the varying intensities of GFP and mCherry in different tissues.

### EM Image Analysis

EM data were quantified using the Image J software. The pixel size of the TEM images was calibrated using a standard sample (diffraction grating replica with latex spheres, TED PELLA, INC, prod. #673) at different magnifications. The values are shown in Table S3. The quantitative data were analyzed by GraphPad Prism 8.4.3.

## ACKNOWLEDGEMENT

We thank Dr. Xiao-Chen Wang of the Institute of Biophysics, Chinese Academy of Science, for providing the anti-VIT-2 antibody; Drs. Cheng-Gang Zhou and Bin Liang, both of Yunnan University, for providing the *ceh-60* mutant, the *vit-2* mutant, and the *vit-2::gfp* knock-in worm strains; and the Caenorhabditis Genetics Center, which is supported by the NIH Office of Infrastructure Programs (P40 OD010440), for providing the wild-type N2 strain. We thank Drs. Wan-Zhong He and Zhao-Di Jiang of National Institute of Biological Sciences (NIBS), Beijing for technical guidance in EM sample preparation. We thank Dr. Jian-Guo Zhang, Dr. Gang Ji, and Can Peng of the Institute of Biophysics, Chinese Academy of Science, for guidance in EM imaging. We are grateful to Dr. John Hugh Snyder for critical reading and editing of this manuscript. This work was funded by National Natural Science Foundation of China (NSFC-ISF 31925026 to F.S., 31501160 to X.-X.L., and 32061143020 to M.-Q.D.), and Ministry of Science and Technology of China (institutional grants to NIBS, Beijing, a fund of the National High-Level Talents Special Support Program to M.-Q.D.), Beijing Municipal Science and Technology Commission (institutional grants to NIBS, Beijing and a fund for cultivation and development of innovation base to M.-Q.D.).

## AUTHOR CONTRIBUTIONS

M.-Q.D. and F.S. supervised the project. M.-Q.D. and C.Z. conceived the project, designed the experiments, interpreted data, and drafted this manuscript. C.Z., N.Z., X.-X.L. and X.-K.T. performed the EM sample preparation. C.Z. and N.Z. constructed worm strains. C.Z. performed electron microscopy imaging, light microscopy imaging, western blotting, and data analysis. All authors read and approved the final manuscript.

## DATA AVAILABILITY STATEMENT

The authors confirm that the data supporting the findings of this study are available within the supplementary material and corresponding authors, upon reasonable request.

## SUPPORTING INFORMATION

| Filename | Description |
| --- | --- |
| Figure_S1 | Quantification of colocalization between VIT-2::GFP and mCherry-tagged VIT-1, -3, and -6 |
| Figure_S2 | VVs and gut granules are two distinct intestinal granules in *C. elegans*. |
| Figure_S3 | Negative controls for the indirect immuno-fluorescent data displayed in Fig. 9 |
| Figure_S4 | YOs detected in -4 -5 and -6 oocytes, albeit fewer in number than in late-stage oocytes from -1 to -3 |
| Figure_S5 | *C. elegans* body wall muscles and pharyngeal muscles failed to be labeled by anti-VIT-2 immuno-gold particles above the background level. |
| Table_S1 | Phosphorylated_site_on_VITs_in_*C. elegans* |
| Table_S2 | Protein-protein_interactions_among_*C. elegans*_VITs |
| Table_S3 | Pixel size in different magnification corrected by standard sample |
| File_S1 | DNA sequence of the homologous templates for *vit-1::mCherry* KI |
| File_S2 | DNA sequence of the homologous templates for *vit-3::mCherry* KI |
| File_S3 | DNA sequence of the homologous templates for *vit-6::mCherry* KI |
| File_S4 | *vit-1* mutant sequence of *vit-1(hq531) vit-2(ok3211)* |
| File_S5 | *vit-1* mutant sequence of *vit-1(hq532) vit-2(ok3211)* |
| File_S6 | *vit-1* mutant sequence of *vit-1(hq533) vit-2(ok3211)* |

---

**Figure S1.**
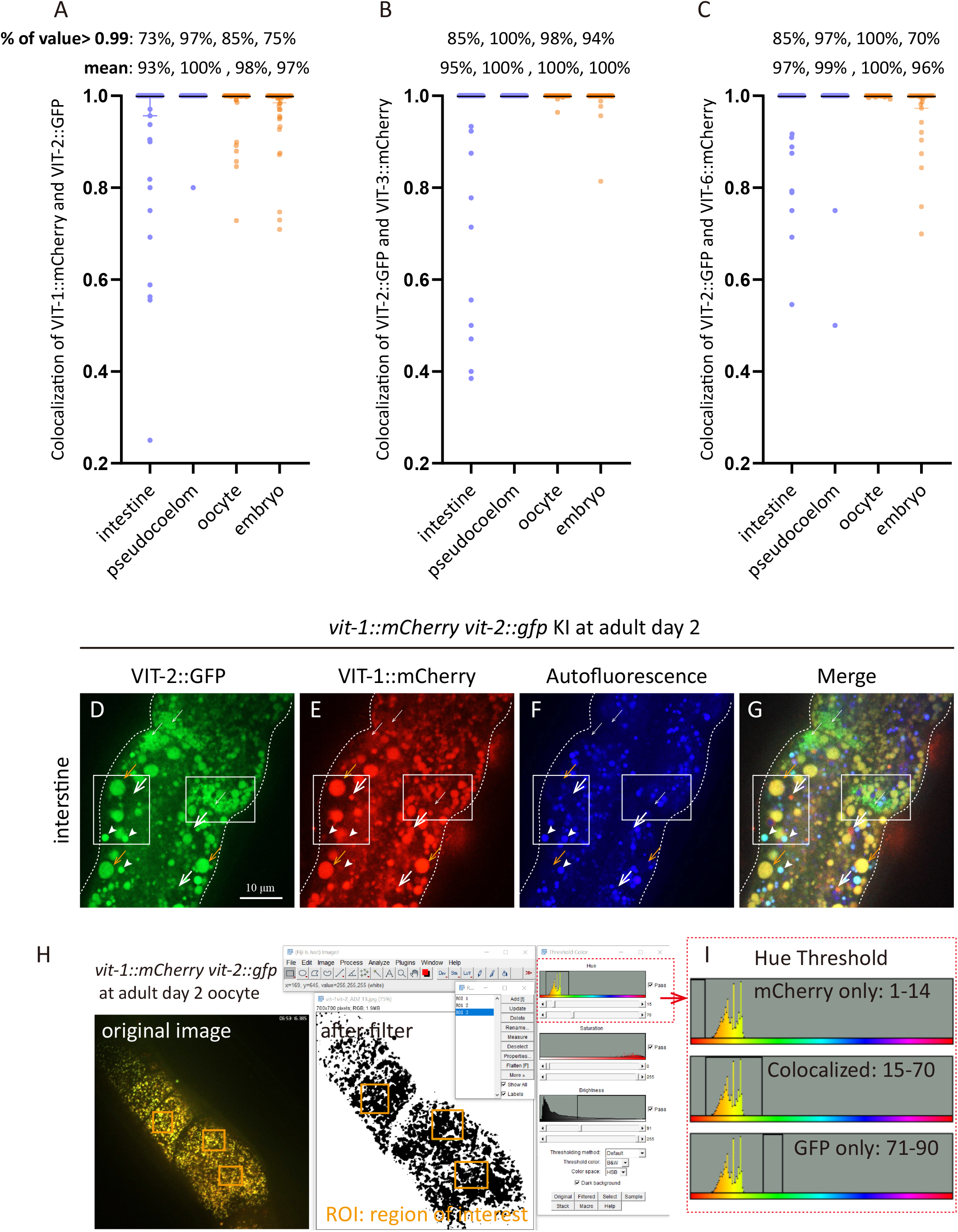
Quantification of colocalization between VIT-2::GFP and mCherry-tagged VIT-1, -3, and -6 (A-C) Tissue-specific quantification results of colocalization between VIT-2::GFP and VIT-1::mCherry (A), VIT-3::mCherry (B), or VIT-6::mChery (C) in young adult worms of AD 1-AD 3. Each point represents one ROI and is color-coded to indicate that the method of quantification-lavender blue for counting the numbers of colocalized and non-colocalized puncta within an ROI or orange for counting the pixels. For the intestine, pseudocoelom, oocyte, and embryo, the numbers of quantified images were 18, 37, 16, and 18 for (A), 22, 46, 20, and 19 for (B), and 20, 70, 14, and 10 for (C). (D-G) show the rare examples of VIT-2::GFP only (thin white arrows) and VIT-1::mCherry only (thick white arrows) VVs. They are tiny compared to VVs containing both VIT-2::GFP and VIT-1::mCherry (orange arrows). Arrowheads indicate autofluorescent gut granules. (H-I) show how colocalization is quantified. See Materials and Methods, Quantification of colocalization between VITs, for details.

**Figure S2.**
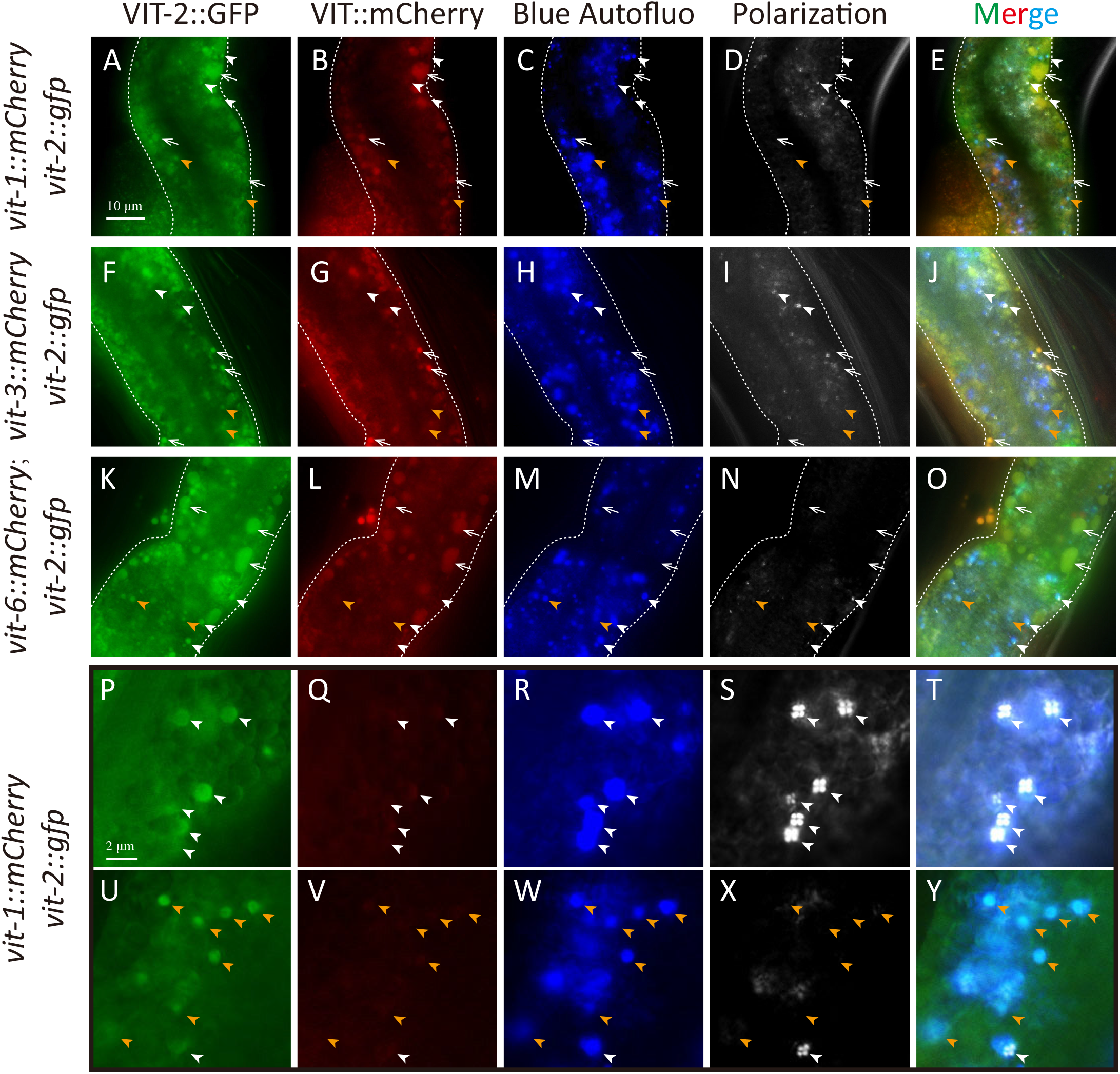
VVs and gut granules are two distinct intestinal granules in *C. elegans*. (A-Y) VVs (white arrows) and autofluorescent gut granules with (white arrowheads) or without (orange arrowheads) birefringent contents in AD 2 worms. (P-Y) Close-up images of gut granules with or without birefringent materials.

**Figure S3.**
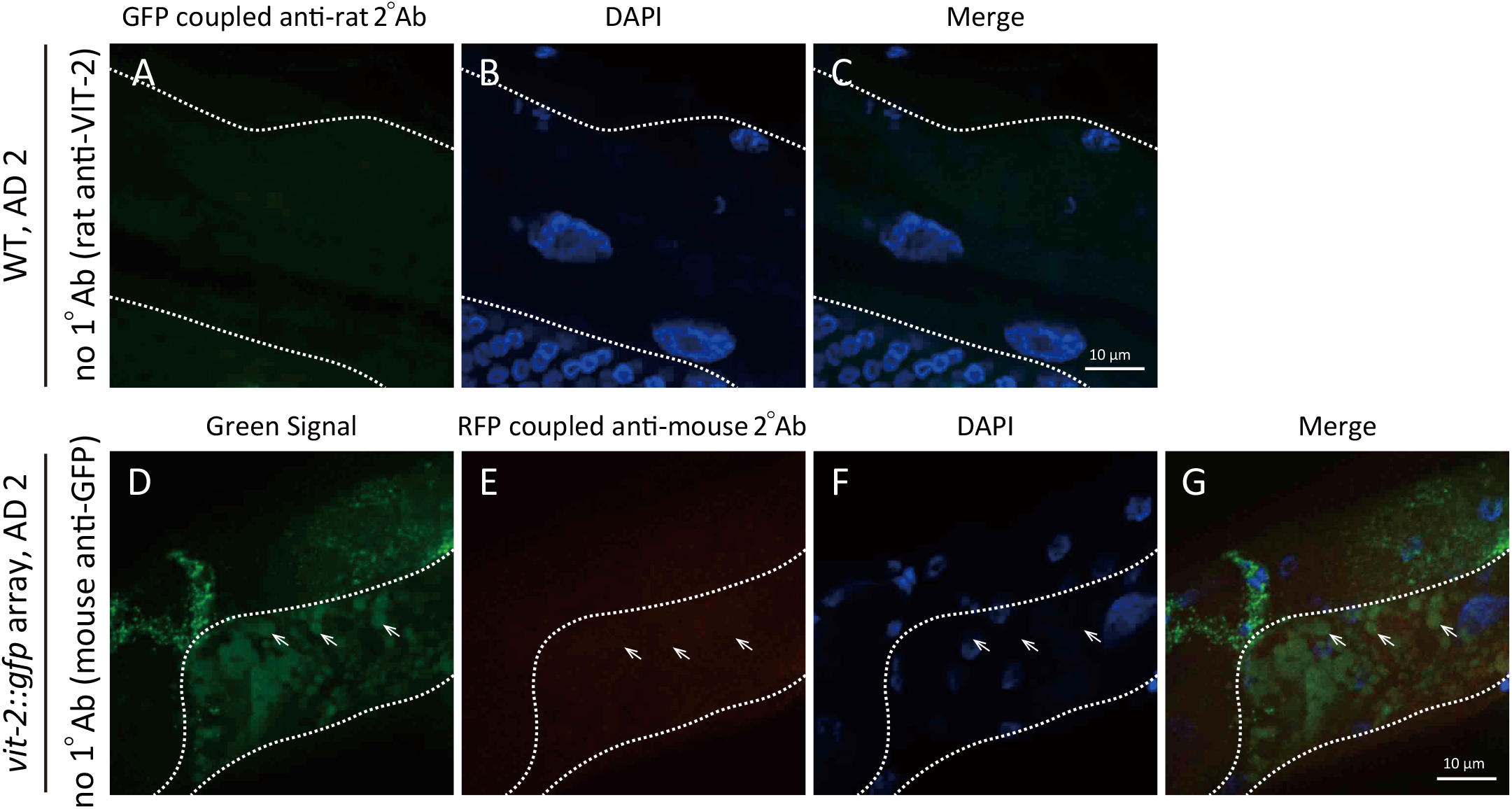
Negative controls for the indirect IF data displayed in figure 9 Primary antibodies were omitted during IF staining, that is, no anti-VIT-2 antibody for (A-C) and no an-GFP antibody for (D-G). White dots outline the intestine. White arrows indicate VVs.

**Figure S4.**
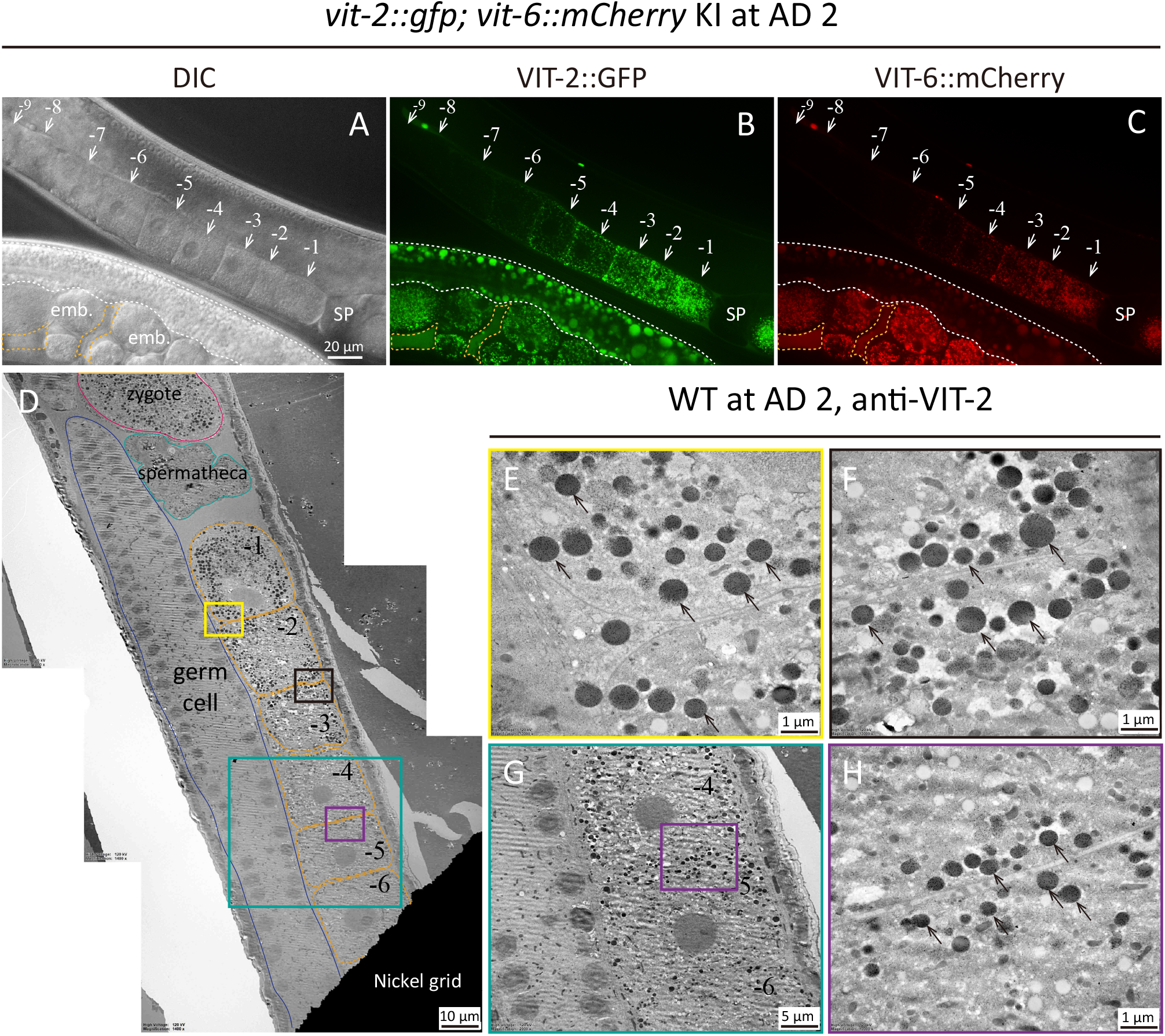
YOs detected in -4, -5, and -6 oocytes, albeit fewer in number than in late-stage oocytes from -1 to -3 (A-C) Light microscopy of the oviduct of an AD 2 WT worm expressing VIT-2::GFP and VIT-6::mCherry, showing DIC (A), green (B), and red (C) fluorescence images. White dots outline the intestine, and orange dots circle the pseudocoelom. White arrows point to oocytes. SP is spermatheca, and emb. means embryo. (D-H) Anti-VIT-2 immuno-EM of WT *C. elegans* on AD 2. Subregions of (D) as marked by the yellow, black, cyan, and purple boxes are shown in (E-H) at higher magnification. Black arrows point to YOs.

**Figure S5.**
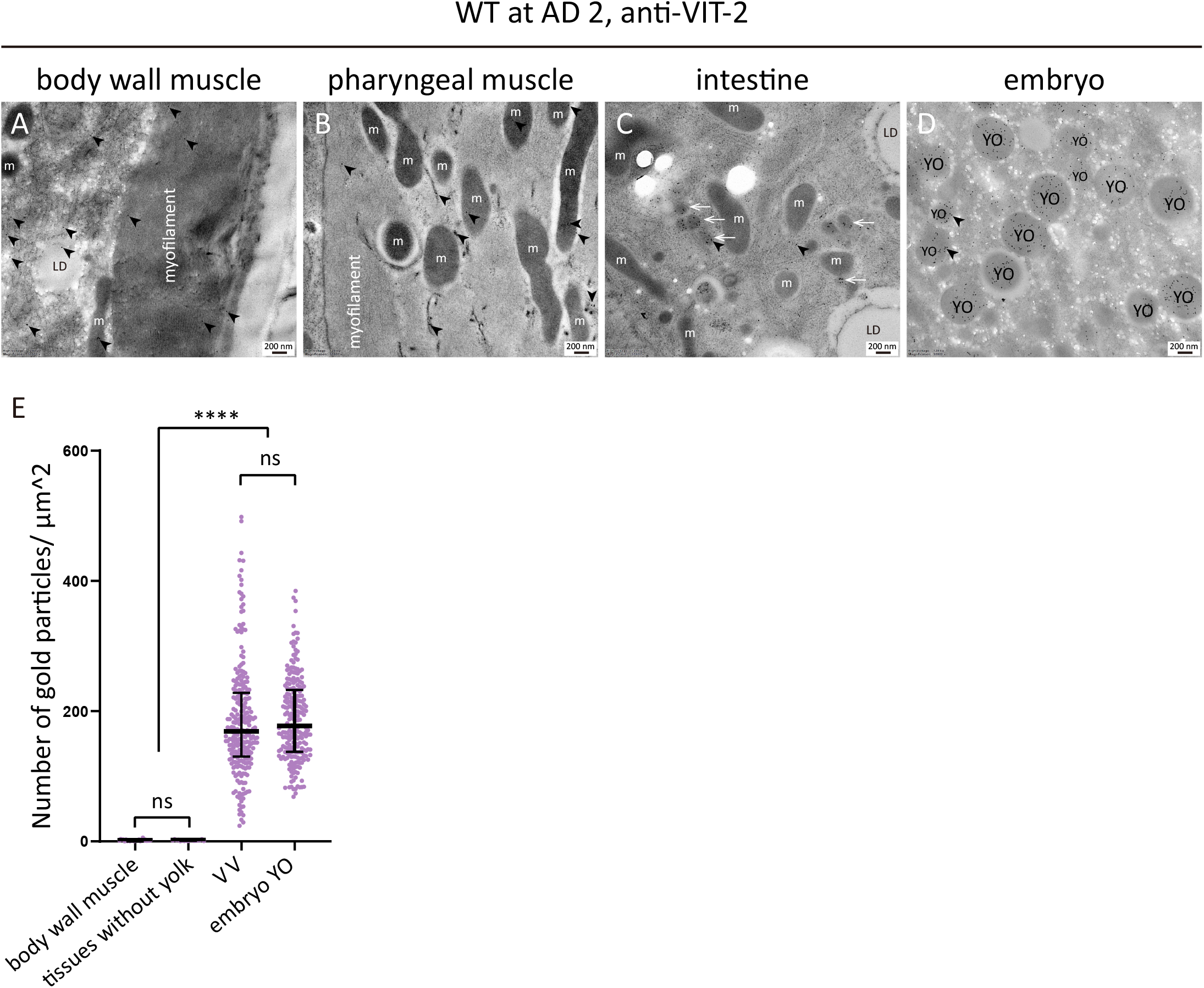
*C. elegans* body wall muscles and pharyngeal muscles failed to be labeled by anti-VIT-2 immuno-gold particles above the background level. Anti-VIT-2 immuno-EM of WT *C. elegans* on AD 2. Representative images are shown for body wall muscles (A), pharyngeal muscles (B), the intestine (C), and embryos (D). White arrows indicate VVs. Black arrowheads point to gold particles. Note that the gold particles in (C) and (D) are numerous and only two are marked by arrowheads in each micrograph, whereas in (A) and (B), all the gold particles are marked. Median and interquartile range are indicated (E). One point represents one value calculated based on one image. 25, 17, 44, and 23 images were quantified for the density of gold particles labeled on body wall muscles, tissues without yolk, VV, and embryo YO. The ssues without yolk include pharyngeal muscles, the hypodermis, the distal gonad, microvilli, and the gonad sheath. \*\*\*\**p* < 0.0001; ns, not significant; one-way ANOVA with Tukey’s multiple comparisons test.

